# Programming cellular condensates for living materials using an intrinsically disordered protein display platform (iDP^2^)

**DOI:** 10.1101/2025.03.02.640994

**Authors:** Rong Chang, Hann Tu, Neel S. Joshi

## Abstract

Self-organization of multicellular systems is vital for building structure in living system but remains underexplored in engineered living materials. We developed iDP^2^, a platform enabling high-density display of intrinsically disordered proteins on the surface of *E. coli* using CsgF as a surface-bound anchor protein to which other protein domains can be fused. Display depends on a lack of intrinsic structure and enable multivalent weak interactions that drive environmental responsiveness and sequence-programmable self-segregation. iDP^2^ allow the programming of material properties like viscoelasticity, stability, offering a method to rationally control cell-cell interactions. This approach could be applied to engineer self-organizing tissues or materials with customized properties.

## Main

Engineered Living Materials (ELMs) is a sub-field that seeks to combine living cells with synthetic systems to create materials with desirable features that mimic natural biology, like self-regeneration, biocatalysts, adhesion, etc^1–3^. A major goal within this field is to control processes like hierarchical structure formation and morphogenesis. Indeed, bio-derived materials like wood, bone, and other tissues have long been a source of inspiration for materials scientists^4^. Only recent advances in synthetic biology have allowed researchers to progress toward programming cells genetically to make such materials through rational design^5,6^. Many early ELM efforts focused on engineering bacteria to produce, secrete, and assemble extracellular biopolymers like cellulose^7^ or proteins to form amyloid fibers (like curli)^5,8^. A less well-studied approach to engineering cells to make materials involves genetically manipulating them to aggregate directly through cell-cell interactions into super-cellular structures. This approach to structure building would be closer to mechanisms of morphogenesis that govern developmental biology of eukaryotes^9,10^. Some progress has been made toward rationally designing such cell-surface interactions to program specific types of cell aggregation. For example, various engineered cellular adhesion molecules (CAMs)^11^ have been used to modify the cell surface and promote self-organization of the cells themselves through a range of protein-protein interactions (such as coiled coils^12^ and nanobody– antigen^13^), with varying levels of specificity.

Existing CAMs have relied almost exclusively on specific molecular recognition to promote cellular aggregation, but this approach hinders the dynamics that facilitate cell reorganization in natural tissue development. Biocondensates, which exhibit dynamic reversibility, could provide an advantage in programmed morphogenesis. Biocondensates occur when biomolecules undergo liquid-liquid phase separation (LLPS) and play an important role in the formation of extracellular matrices (e.g., bio-adhesives^14,15^, tissue tight junctions^16^), and intracellular membrane-less organelles (i.e., via the condensation of proteins, RNA, and other biomolecules)^17^. Many biocondensates, such as stress granules and p-bodies^18^ rely on intrinsically disordered proteins (IDPs), whose high conformational flexibility and multivalent weak interactions drive phase separation^19,20^ (Fig. 1a). Condensate-forming IDPs offer the attractive opportunities to programmed morphogenesis and ELMs to include new controllable driving forces for structure building based on phase separation^21^. To date, the only reported case of ELM formation with IDP aggregation as a driving force involved elastin-like peptides (ELP). These were displayed on the surface of *Caulobacter crescentus*^22^ or *E. coli*^23^ to create a system that spontaneously forms a self-standing material during microbial cell culture. However, the full scope of IDP sequence space is worthy of exploration, as it includes various homotypic and heterotypic interactions with different specificities. Using IDPs as unique CAMs (Fig. 1b) will also require well-characterized surface-display mechanisms with high surface density.

**Fig. 1.**
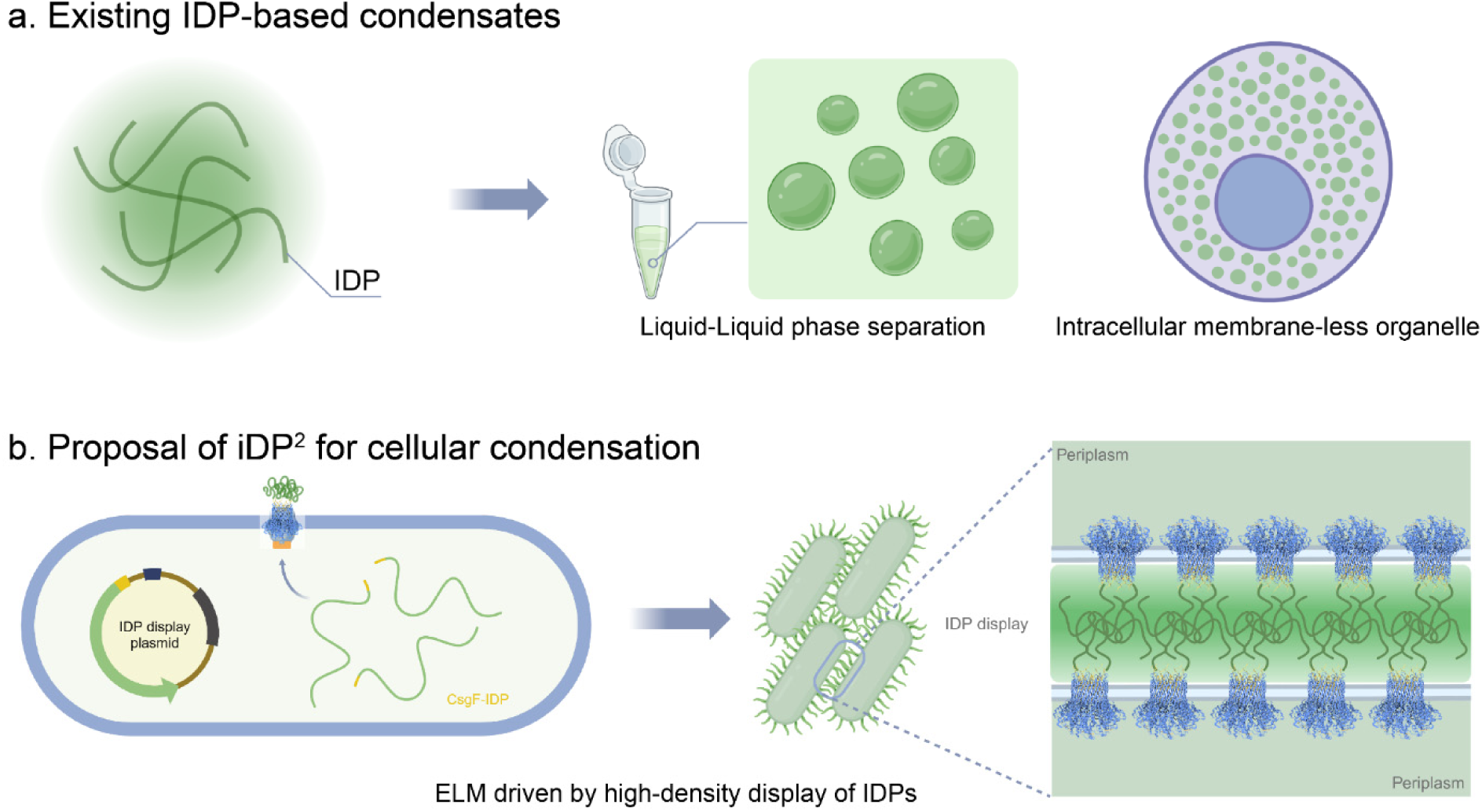
**a**, Existing IDP-based condensates including liquid-liquid phase separation in solution and intracellular membrane-less organelle formation. **b**, Proposal for iDP^2^ for cellular condensation in this work, which drives ELM formation using high-density display of IDPs.

Here we demonstrate new way to repurpose the well-studied curli (i.e., Type VIII) secretion system as a high-density cell surface display pathway in *E. coli* for the purpose of promoting cellular aggregation through biocondensate formation (iDP^2^). This pathway uses dedicated outer-membrane secretion machinery that relies on an entropy-driven mechanism to secrete several proteins from the curli operon (e.g., CsgA, CsgB, CsgF). (Fig. 2a-c) We demonstrate for the first time that CsgF can be used as an anchor for very high-density display on the outer membrane of *E. coli*. iDP^2^ demonstrates that IDPs can direct whole-cell assembly into distinct patterns with a high degree of sequence selectivity, stimulus responsiveness, and straightforward protocols. The resulting cellular aggregates show promise for ELM production based on the tunability of their materials properties (e.g., viscoelasticity, stability, spatial patterning) by appropriate choice of IDP sequence.

**Fig. 2.**
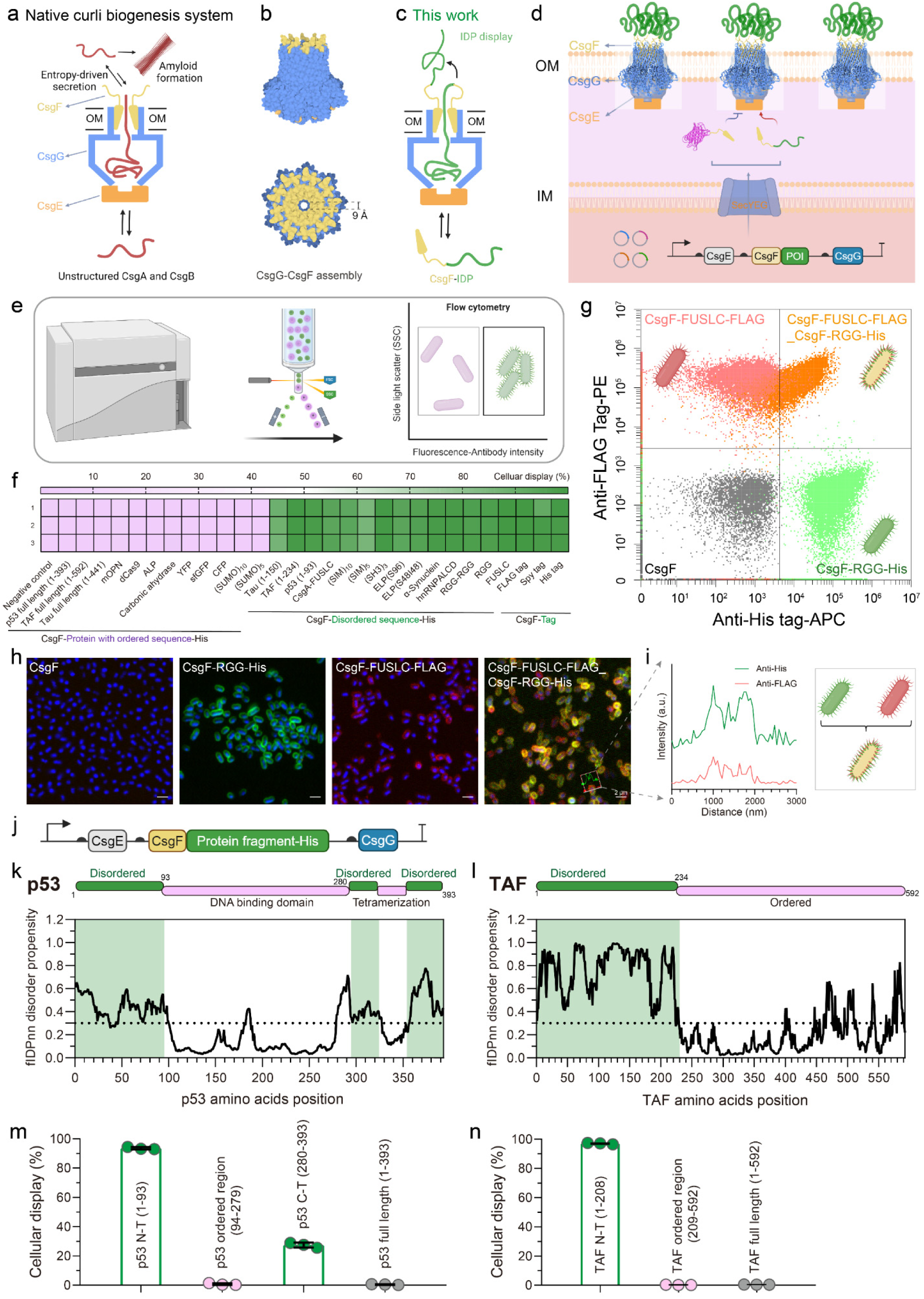
The iDP^2^ platform displays intrinsically disordered proteins on the surface of *E. coli*. **a**, Schematic of outer membrane (OM) pore complex of curli biogenesis system, composed of CsgE-CsgG-CsgF, and dedicated to the secretion of amyloidogenic CsgA and CsgB. **b**, CsgG-CsgF complex crystal structures from Protein Data Bank; blue is CsgG and yellow is CsgF (PDB code: 6L7A). **c-d**, Schematic of CsgF-IDP fusion and surface display. **e**, Flow cytometry workflow to screen the various protein sequences fused to CsgF at the C-terminus. **f**, Display efficiency of various CsgF fusions, including structured, unstructured, and affinity tag domains. **g-h**, Flow cytometry (**g**), and CLSM images (**h**) of immunostained PQN4 displaying representative IDPs including FUSLC, RGG, or both FUSLC and RGG together. FUSLC is labeled with a His tag (green) and RGG with a FLAG tag (red); all cells are labeled with the nucleic acid stain DAPI (blue). CsgF display with no IDP or affinity tag is shown as a negative control. Scale bars of (**h**): 2 μm. **i**, Fluorescence intensity profiles across cells displaying both FUSLC and RGG presented in (**h**) and scheme for multiple-IDP display. **j**, Genetic construct design for CsgF fused with various protein fragments, co-transcribed with CsgE and CsgG. **k-l**, Annotated protein domain maps (from DisProt database^33^) and disorder prediction as a function of amino acid position (by flDPnn^34^) for p53 (**k**) and TAF (**l**). Disordered domains are highlighted in green and ordered domains in blue. **m-n**, Display efficiency of CsgF fused with either the whole sequence or truncated domains of p53 (**m**) and TAF (**n**) as measured by flow cytometry. Cellular display data of (**f**) and (**m-n**) represents the percent of cells above a threshold determined by CsgF-His strains serving as positive control and CsgF only strains as negative control. Threshold position is shown in Extended Data Fig. 1a-c.

### Design and validation of IDP display using CsgF

The Type VIII secretion system is to be compatible primarily with unstructured proteins. Its native secretion targets – known CsgA and CsgB – are thought to be kept in a disordered state by the chaperone, CsgC, inside the cell to facilitate translocation through the CsgG porin. (Fig. 2a) CsgE and CsgF are thought to bind to different faces of CsgG^24^ (Fig. 2a). On the periplasmic side, the CsgE-CsgG complex captures unfolded proteins based on recognition of a N-terminal amino acid sequence and confines them, creating an entropically-driven free-energy gradient promoting translocation outside the cell^25^. On the extracellular face, the CsgG-CsgF complex forms a 9 Å diameter channel that only accommodates proteins in extended, unfolded conformations^26^ (Fig. 2b). We rationalized that the native function of the Type VIII machinery makes it ideal for IDP secretion and display.

We demonstrated this by creating a library of 26 different CsgF fusion proteins^27^, where the domains fused to the C-terminus varied in their degree of inherent structure. We expected that, after secretion through CsgG, the CsgF fusion proteins would be displayed on the cell surface. (Fig. 2c) The CsgF chimeras were constituted in a plasmid library where each chimera was co-transcribed with genes encoding CsgE and CsgG. (Fig. 2d) The CsgF chimeras all contained a C-terminal affinity tag for detection via antibody labeling. Each plasmid library member was transformed into a laboratory strain of *E. coli* (PQN4) that has the native curli genes deleted from the chromosome^28^. Screening of the library by flow cytometry after induction in liquid culture and antibody labeling revealed a clear preference for unstructured over structured domains (Fig. 2e-f, Extended Data Fig. 1a-c and 2a). Two IDPs in particular – the low-complexity domain of FUS RNA binding protein (FUSLC)^29^ and the arginine/glycine-rich domain from the P granule protein LAF-1 (RGG)^30^ – showed a high degree of cell surface display based on flow cytometry (Fig. 2g), immunostained fluorescence microscopy (Fig. 2h and Extended Data Fig. 2b), and western-blot (Extended Data Fig. 2c). Compared with the amount of CsgF-IDPs in the cellular fraction, there was a small amount of CsgF-IDP found in the medium, confirming that most of the secreted protein was captured by the cell (Extended Data Fig. 2c). Both FUSLC and RGG could even be displayed on the surface of the same cell transformed with a plasmid encoding genes for both CsgF-FUSLC and CsgF-RGG (Fig. 2g-i and Extended Data Fig. 2b-c), suggesting that this could be a generalizable approach to displaying multiple IDP sequences simultaneously.

To further confirm that the CsgF-IDP chimeras were being secreted through the curli biosynthesis machinery, we monitored their cell surface display in strains with and without CsgC and CsgG. Flow cytometry showed that cells missing CsgG did not exhibit fluorescence signal as expected, since CsgG forms the outer membrane pore. In comparison, CsgC, seemed to be less important for CsgF-IDP display. This is in line with CsgC’s expected role of preventing folding of the native curli substrates (CsgA and CsgB) inside the cell^24^ (Extended Data Fig. 2d).

### Use of iDP^2^ to distinguish between structured and unstructured protein domains

Many naturally occurring proteins contain both structured and unstructured domains within their sequence. To confirm that iDP^2^ was exclusively compatible with only unstructured domains, we considered two proteins (p53 and TAF)^31^ that exhibit a mix of order and disorder in their structure. Fig. 2j shows their degree of order as a function of amino acid position, as predicted by bioinformatics tools from DisProt, a database dedicated to cataloguing IDP structure and function^32^ (Fig. 2k-l). When either the full-length protein sequences or only the structured domains were fused to CsgF, we saw no evidence of surface display based on flow cytometry. However, when only the unstructured domains of both proteins were fused to CsgF, we observed robust surface display (Fig. 2m-n). This may suggest that iDP^2^ could be useful as a tool to distinguish between domains with and without structure while avoiding more labor-intensive conventional structural biology approaches.

### Quantification of surface display density for iDP^2^

Since biocondensate formation is highly dependent on concentration, we thought it would be important to have high surface display density to promote cellular aggregation and initiate ELM formation. Therefore, we quantified surface display density with both western-blot and flow cytometry. First, we fractionated the various extracellular and sub-cellular compartments – secreted, membrane, periplasmic, cytosolic, and inclusion bodies – after induction of CsgF chimeras and subjected them to analysis by western blot^35^ (Fig. 3a). We found that CsgF-His (5.0%) and a representative IDP fusion (CsgF-FUSLC-His, 4.4%) exhibited a similar percentage of total protein localized to the surface (i.e., membrane fraction). (Extended Data Fig. 3a-c) Most of the overproduced proteins were relegated to the inclusion body and cytosolic fractions (Extended Data Fig. 3a-c). We used a calibration curve of known CsgF-His concentrations to estimate the absolute concentration of protein in the membrane fraction from western blot band intensity (Fig. 3b and Extended Data Fig. 3d-e). This, combined with CFU counting (Extended Data Fig. 3f-g) prior to cell lysis, allowed us to calculate the number of surface-displayed CsgF-His per cell surface to be (16.1 ± 4.9) × 10^4^ and the mean surface density to be (3.6 ± 1.1) × 10^4^ µm^-2^ (Fig. 3e, Table 1, Note S1). To corroborate the results obtained by western blot, we also generated a calibration curve for fluorescence intensity using beads (Ultra Rainbow Fluorescent Beads) with known fluorophore surface density (Fig. 3c-d). This technique led to an estimate of (7.4 ± 0.4) × 10^4^ CsgF-His per cell and a surface density of (1.6 ± 0.1) × 10^4^ µm^-2^ (Fig. 3e, Table 1, Note S1).

**Fig. 3.**
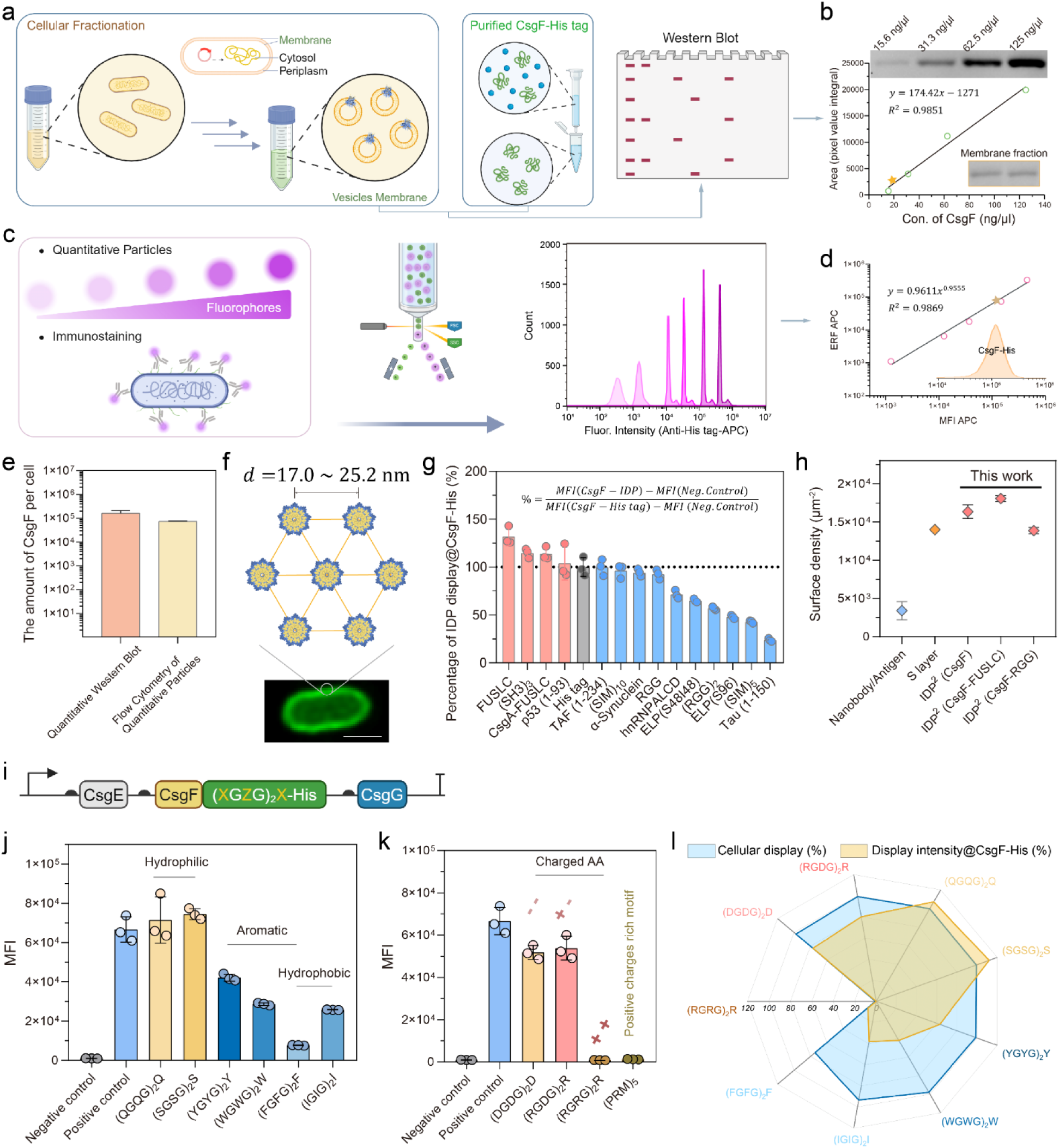
Quantification of the surface display density of iDP^2^ and its sequence heuristics. **a**, Schematic of cellular fractionation and purification of CsgF-His for quantitative western blot. **b**, The fitted standard curve of quantitative western blot via its pixel value integral of purified CsgF-His tag with different concentrations and membrane fraction samples. **c**, Schematic of polymer microparticles with known amounts of fluorophore immobilized and cell surface immunostaining with an antibody bearing the same fluorophore. **d**, The fitted standard curve of standard fluorescently labeled particles via equivalent number of reference fluorophores (ERF) and mean fluorescence intensity (MFI). Fluorescence intensity histogram of CsgF-His strains shown in inset. The mean values for the positive control sample (CsgF-His) were indicated with ⋆ in (**b**) and (**d**). **e**, The calculated amount of CsgF per cell based on quantitative western bot and flow cytometry of standard particles. **f**, Schematic of CsgF-CsgG complexes with calculated distance between each complex, assuming even distribution on the cell surface. Scale bars: 1 μm. **g**, Mean intensity of surface display for various CsgF-fusion constructs, with CsgF-His normalized to 100%. **h**, Comparison between the density of cell surface display for iDP^2^ and other published systems^22,36,37^. **i**, Genetic constructs design for CsgF fused de novo sequences, where X and Z are variable amino acids. **j**, Display intensity of CsgF-(XGZG)_2_X based on MFI values from flow cytometry. **k**, Flow cytometry-based display intensity of CsgF fused to domains with different electrostatic charge properties: neutral zwitterionic – (RGDG)_2_R; negative – (DGDG)_2_R; and, positive – (RGRG)_2_R and (PRM)_5_. **l**, Radar map showing percentage of cellular display and display intensity compared to CsgF-His tag for all designed sequences.

**Table 1.** Quantification of surface density of iDP^2^.

| CsgF-IDP | Quantitative Western Blot |  |  | Quantitative Flow Cytometry |  |  |
| --- | --- | --- | --- | --- | --- | --- |
| | # of proteins per cell surface | Surface density ($\mu\text{m}^{-2}$) | Averaged distance between adjacent curli channels (nm) | # of proteins per cell surface | Surface density ($\mu\text{m}^{-2}$) | Averaged distance between adjacent curli channels (nm) |
| CsgF | $(16.1 \pm 4.9) \times 10^4$ | $(35.7 \pm 11.0) \times 10^3$ | $17.0 \pm 2.6$ | $(7.4 \pm 0.4) \times 10^4$ | $(16.3 \pm 0.9) \times 10^3$ | $25.2 \pm 0.7$ |
| CsgF-FUSLC | NA | NA | NA | $(8.1 \pm 0.2) \times 10^4$ | $(18.1 \pm 0.4) \times 10^3$ | $24.0 \pm 0.3$ |
| CsgF-RGG | NA | NA | NA | $(6.2 \pm 0.2) \times 10^4$ | $(13.9 \pm 0.4) \times 10^3$ | $27.4 \pm 0.4$ |

We considered that our estimates of cell surface density of CsgF fusion display may include systematic error. We rationalized that western band intensity could be an overestimate if it included elements of the inner membrane and periplasm. On the other hand, steric hindrance or other labeling inefficiencies could cause the surface-bound antibodies on the cell to underestimate the total CsgF on the surface during flow cytometry. Taking these factors into account, we estimated that the amount of CsgF-His per cell to fall within the range of 7.4 x 10^4^ - 16.1 × 10^4^, which is higher than most published systems. Furthermore, if we assume that the CsgF fusions are uniformly distributed across the cell surface, we calculated that the average distance between two CsgG-CsgF complexes on the cellular surface was between 17.0 - 25.2 nm^25^ (Fig. 3f). Since the formation of the outer membrane pore complex involves the assembly of 9 CsgG-CsgF monomers we could estimate an even higher surface density of CsgF fusions (Table 1). Using CsgF-His as the benchmark, we compared the display density of all the CsgF-IDP fusions on a relative basis (Fig. 3g). Most of the CsgF-IDPs displayed with comparable density to CsgF-His and some were even higher than other published high-density display systems^22,36^ (Fig. 3h). Overall, there was a general trend showing that the mean number of proteins per cell surface decreased with higher length protein sequences fused to CsgF (Extended Data Fig. 3h-i).

### Sequence heuristics of iDP^2^

After establishing the high surface density of iDP^2^ and its compatibility with a wide range of unstructured sequences, we sought a more systematic understanding of sequence effects the fused domains. We designed and created a library of short, de novo fusion peptides with the sequence (XGZG)_2_, where X and Z could be any standard amino acid and the high glycine (G) frequency would ensure a disordered domain (Fig 3i). In almost all cases where X and Z were identical, surface display was observed to varying degrees by flow cytometry analysis (Extended Data Fig. 3j). Uncharged hydrophilic amino acids – (SGSG)_2_S and (QGQG)_2_Q – performed comparably to CsgF-His, presumably because they maintained a flexible and hydrated conformation during secretion (Fig. 3j). As for aromatic and hydrophobic residues (Y, W, F, I), none showed more than 50% surface density compared with CsgF-His (Fig. 3j). We speculate that the steric hindrance of bulkier side chains and potential for hydrophobic interactions with the membrane led to less surface display.

Negatively charged ((DGDG)_2_D) and neutral zwitterionic ((RGDG)_2_R) fusion domains showed >99% cellular display and ∼80% surface density compared CsgF-His (Fig. 3k and Extended Data Fig. 3k). The only sequence for which no cell surface display was observed was (RGRG)_2_R. We speculate that the highly positively charged sequence interfered with membrane integrity, which was corroborated by the failure of another positive charge-rich motif, (PRM)_5_ (Fig. 3k and Extended Data Fig. 3k). Fig. 3l summarizes the heuristic results: small, hydrophilic, or negatively charged sequences were most compatible with iDP^2^, while bulky, hydrophobic, or positively charged sequences hindered or abolished cell surface display.

### iDP^2^ for cellular aggregation driven by multivalent weak interactions

Phase separation of IDPs through multivalent weak interactions is known to occur in various biological contexts. Since iDP^2^ led to very high density IDP surface display, we sought to leverage this to promote cell condensation and self-organization. Accordingly, we subjected 12 strains displaying different IDPs to an aggregation assay. The assay was based on sedimentation under static conditions in tubes as measured by OD_600_ in the upper half of the tubes after overnight incubation at different temperatures (Fig. 4a). Strains displaying some of the IDPs (FUSLC, RGG, (RGDG)_2_R, hnRNPA1LCD, TAF (1-234), and ELP) enhanced sedimentation rate, which we took as evidence of potential aggregation driven by IDP phase separation (Fig. 4b). Indeed, aggregation and settling could be observed in some tubes with the naked eye even after standing for only 1-2 hours, while the control group showed unchanged turbidity (Fig. 4a). CsgF-His alone exhibited low aggregation propensity at all three temperatures. Four strains displaying IDPs (FUSLC, RGG, (RGDG)_2_R, hnRNPA1LCD) showed ∼90% aggregation propensity at all temperatures (Fig. 4c).

**Fig. 4.**
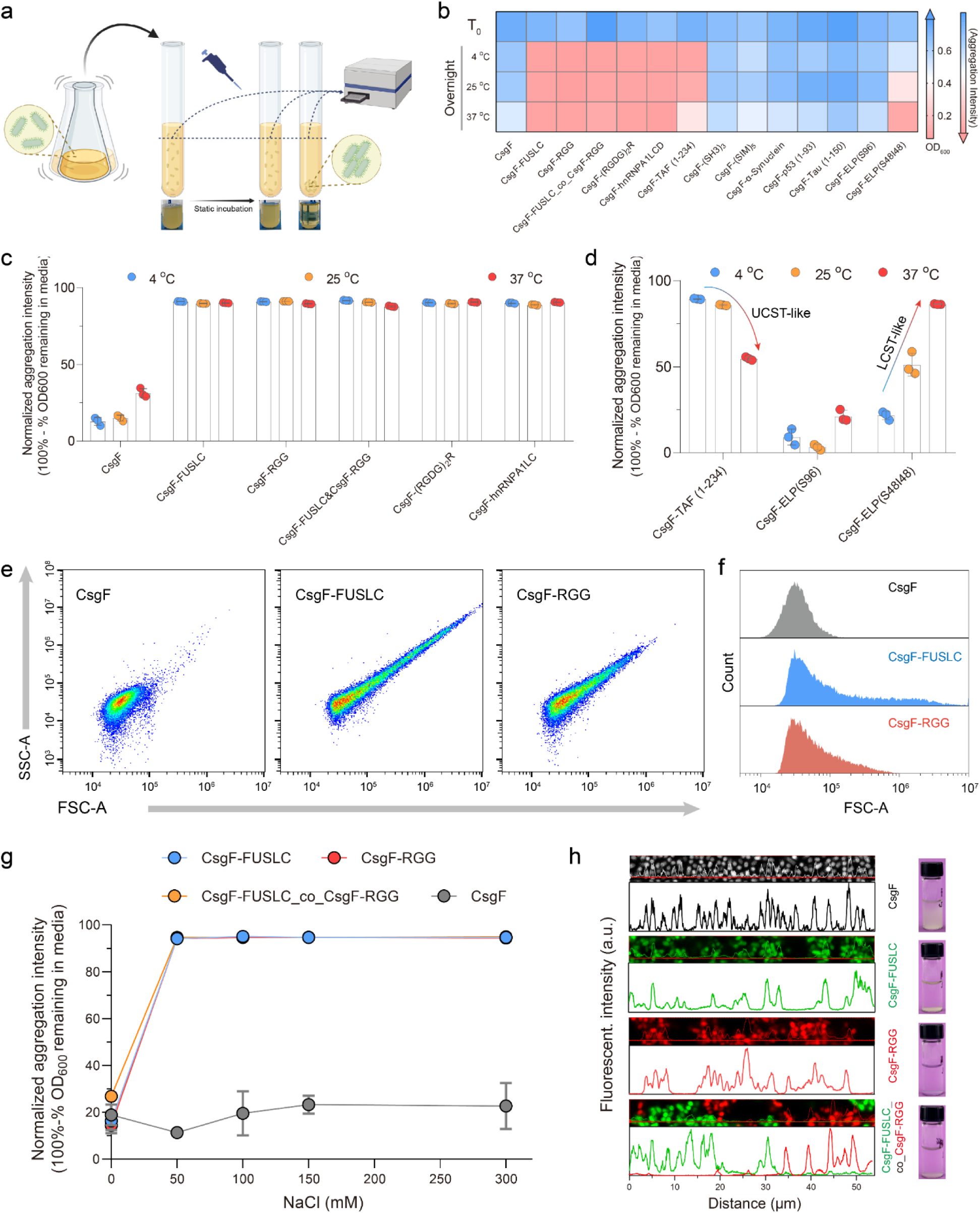
Bacterial aggregation via surface display of IDP with multivalent weak interaction. **a**, Schematic of aggregation assay for PQN4 strains expressing CsgF-IDP fusions on their surface using OD_600_ measurements. Bottom images show representative sedimentation observable by naked eyes after 1∼2 hours. **b**, Heat maps representing OD_600_ measurements of all PQN4 strains displaying different IDPs. Red indicates lower OD_600_ remaining in suspension, i.e., higher aggregation propensity. Blue indicates the opposite. **c**, Normalized aggregation propensity of bacteria displaying various IDPs after overnight incubation at different temperatures. **d**, Normalized aggregation propensity of bacteria displaying temperature-responsive IDPs after overnight incubation at various temperatures. **e**, Representative flow cytometry dot plots of FSC-A versus SSC-A for PQN4 displaying CsgF, CsgF-FUSLC and CsgF-RGG (n = 5,000). **f**, Corresponding plots of counts as a function of FSC-A values for distinct strains. **g**, Normalized aggregation propensity of bacteria in different concentrations of NaCl, as measured by OD600. Data in **c**, **d**, and **g** normalized to turbidity value before each incubation with fully suspended medium. **h**, Fluorescence microscopy images for strains displaying CsgF and CsgF-IDPs. Fluorescence intensity plots were obtained by cross sectional analysis. Images show sedimentation or turbidity visible by eye.

We also sought to investigate dynamic aspects of IDP-driven aggregation with domains that exhibit known temperature-responsive behaviors. When TAF (1-234), the disordered fragment of a DNA binding protein, was displayed using iDP^2^, cells preferentially aggregated at 4 °C compared to higher temperatures (Fig. 4d). We also fused two different elastin-like peptide (ELP) sequences to CsgF. Each ELP consisted of 96 repeats of the sequence (VPGXG). One variant (S96), in which the X position was occupied by serine, was known to not exhibit temperature-responsive behavior because it was too hydrophilic to undergo phase separation. The other variant (S48I48) was known to exhibit lower critical solution temperature (LCST) behavior^38^. This temperature-responsiveness was preserved in iDP^2^. Strains displaying CsgF-ELP(S96) fusions showed low aggregation propensity regardless of temperature, while those displaying CsgF-ELP(S48I48) preferentially aggregated at 37 °C compared to lower temperatures (Fig. 4d).

Cellular aggregation based on iDP^2^ could also be monitored by light-scattered area (FSC-A) versus side-light-scattered area (SSC-A) dot plots from flow cytometry, where higher SSC-A and FSC-A values indicate cell aggregation. Both CsgF-FUSLC and CsgF-RGG strains showed higher and wider distribution of SSC and FSC than strains without IDPs displayed (Fig. 4e-f). In exploring various conditions to control aggregation state, we unexpectedly discovered that the CsgF-FUSLC/CsgF-RGG pair exhibits responsiveness to salt concentration, with higher NaCl concentrations leading to high aggregation propensities (Fig. 4g), while CsgF alone without IDP fusions does not exhibit this behavior. These results were corroborated by cross sectional analysis of fluorescence microscopy images, which showed intensity spikes corresponding to single cell widths for CsgF, but wider spikes for CsgF-IDP fusions (Fig. 4h).

### Self-patterning and dynamic bacterial aggregation via iDP^2^

Having demonstrated the high-density surface display of iDP^2^ and its ability to direct cellular aggregation, we explored its relevance for multicellular patterning. FUSLC and RGG were compatible with iDP^2^ and the soluble proteins were previously known to undergo orthogonal phase separation – i.e., they would not mix, but would form their own homogenous phases in aqueous solutions under proper conditions^21^. We tried to leverage this capability to direct cell patterning. Accordingly, we created all permutations of PQN4 strains that displayed either FUSLC, RGG, or CsgF only and produced either cytosolic CFP or RFP. We expected these strains in different combinations to exhibit either homogeneous, heterogeneous, or no aggregation (Fig. 5a-c).

**Fig. 5.**
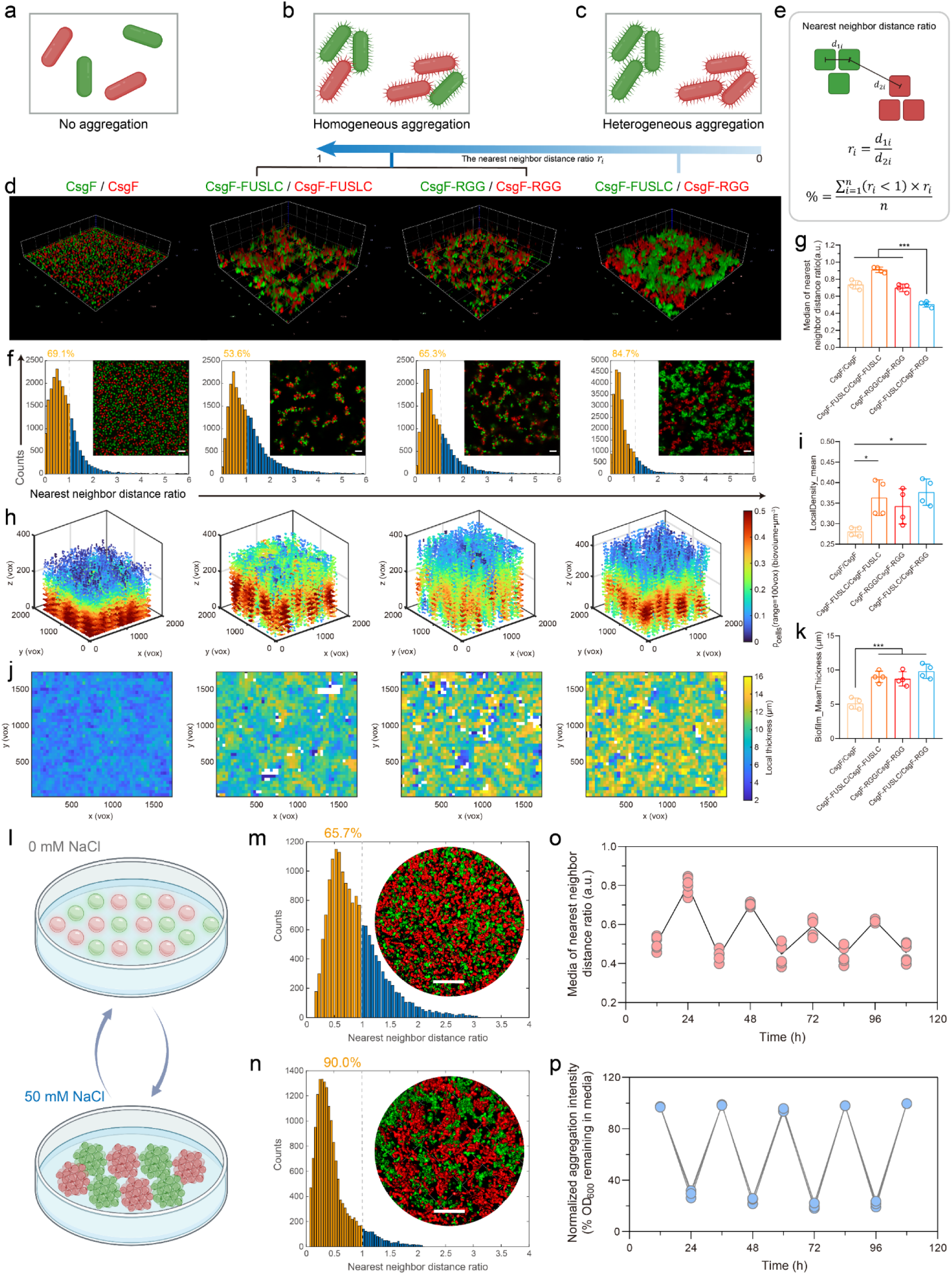
Self-patterned bacterial aggregation via surface display of IDPs with phase separation capability. **a-c**, Schematic explaining different pattern formation when mixing two different strains together including no aggregation (**a**), homogeneous aggregation (**b**) and heterogeneous aggregation (**c**). **d**, CLSM Z-stack of CsgF/CgF, CsgF-FUSLC/CsgF-FUSLC, CsgF-RGG/CsgF-FUSLC and CsgF-FUSLC/CsgF-RGG strain combinations with each strain producing either CFP (green) or RFP (red). **e**, Mathematical explanation of nearest neighbor distance ratio (*r*_i_) analysi used to quantify the cellular pattern formation based on homogenous interactions. *r*_i_ = 1 indicates complete no neighbor preference (i.e., homogenous distribution); 0 < *r*_i_ < 1 indicates a preference to like neighbors (i.e., heterogeneous distribution). Images were segmented uniformly as 1.2 μm cubes (approximating the size of an *E. coli* cell). **f-g** Histogram plots (**f**) and statistical analysis of median values (**g**) of nearest neighbor distance ratio and embedded representative 2D CLSM images of CsgF/CgF, CsgF-FUSLC/CsgF-FUSLC, CsgF-RGG/CsgF-RGG and CsgF-FUSLC/CsgF-RGG combinations. The percentage of segmented cubes satisfying the condition *r*_i_ < 1 (yellow bars) is shown above each histogram. Scale bars: 5 μm. **h-i**, 3D Heatmap (**h**) and statistical analysis (**i**) of bacterial local density for the different cell strain combinations. **j-k**, 2D heatmaps (**j**) and statistical analysis (**k**) of bacterial local thickness for the different cell strain combinations. **l**, Schematic of protocol for dynamic reconfiguration of cellular patterns via resuspension in different NaCl concentrations. **m-n**, Representative histogram plots of nearest neighbor distance ratio for CsgF-FUSLC/CsgF-RGG in 0 mM NaCl (**m**) and 50 mM NaCl (**n**). Scale bars of (**m-n**): 10 μm. **o-p**, Oscillation of median nearest neighbor distance ratio (**o**) and aggregation propensity (**p**) during repeated cycles of resuspension in different salt conditions. P values were determined by one-way ANOVA with Tukey’s post-test; *P < 0.01, **P < 0.01, ***P < 0.001, (n=4 repeats per group).

When different combinations of strains were co-incubated in well plates and allowed to sediment, they produced different patterns, as monitored by confocal fluorescence microscopy (Fig. 5d). When CFP-producing (green) and RFP-producing (red) strains displaying only CsgF were combined, the cells appeared homogeneously distributed with no apparent pattern formation. Bacteria displaying the same IDP (either FUSLC or RGG) on their surface led to larger cellular aggregates, but a homogenous distribution of green and red cells in both cases. In contrast, mixtures of green cells displaying CsgF-FUSLC and red cells displaying CsgF-RGG showed distinctly heterogeneous patterning with larger islands of segregated red and green cells.

We used Biofilm Q to quantify local density and biofilm thickness, and nearest-neighbor analysis to quantify differences in cell patterning for these different strain combinations^39^. The nearest neighbor distance ratio (*r*_i_) was defined as the ratio of distances between the closest in-phase neighbor and the closest out-of-phase neighbor for each cell (Fig. 5e). *r*_i_approaches 1 for completely homogeneous aggregation – i.e., no preference for neighbor identity. The more strains prefer interacting with neighbors that match their own identity, the more the *r*_i_ decreases toward 0. Using Biofilm Q, we segmented the microscopy images into cubes of length of 1.2 μm and generated histograms of *r*_i_ for all strain combinations (Fig. 5f). The fraction of cubes exhibiting *r*_i_ < 1 was signficantly lower for the CsgF-FUSLC/CsgF-RGG combination, compared to the other conditions (Fig. 5f). The median of *r*_i_ for the CsgF-FUSLC/CsgF-RGG mixture (∼0.50) was significantly lower than the other groups (Fig. 5g). Other forms of quantitative and statistical analyses supported higher local cell density and mean biofilm thickness for the homogeneous and heterogeneous combinations compared to CsgF alone (Fig. 5h-k). Strains mixtures displaying other IDP combinations also exhibited interesting patterning features, suggesting that more sequence programmability of IDP-based patterning may be possible (Extended Data Fig. 4a).

A feature of cellular patterning driven by the multivalent weak interactions of IDPs, as compared to other CAM strategies^12,13^, is their dynamic reversibility. Based on the responsiveness to ionic concentration we identified for CsgF-FUSLC and CsgF-RGG (Fig. 4h), we explored the ability to dynamically reconfigure cellular patterns based on solution-phase conditions. This combination could be repeatedly cycled between aggregated and non-aggregated states based on resuspending the cells in 0 or 50 mM NaCl over 4 cycles, as measured by nearest-neighbor analysis from microscopy images (Fig. 5l-p). We did observe some decay in the amplitude of the oscillating signal, which we speculate was related to protein degradation and decreased cell viability over the course of the multi-day cycling experiment. Pattern formation in developmental biology usually requires high cell densities with multiple cell types. We tried to mimic this morphology by combining three iDP^2^ strains: one displaying CsgF-RGG only, one displaying CsgF-FUSLC only, and a third displaying both CsgF-RGG and CsgF-FUSLC simultaneously. The three strains were also labeled through cytosolic expression of a unique fluorescent proteins. We found that at sufficiently high cell densities, the strains expressing CsgF-RGG only and CsgF-FUSLC only auto-segregated, as expected, but when the hybrid strain expressing both IDPs was included, it occupied the interstitial space between the other two cell population, like a border (Extended Data Fig. 4b-d).

### Exploring the compatibility of iDP^2^ with other ELM scaffolds

Although iDP^2^ represents a unique surface display approach for IDPs and exhibits some desirable features (e.g., dynamic reconfigurability), we were also interested in combining it with more mechanically robust ELM scaffolding schemes. Naturally occurring biological tissues usually rely on a mixture of cell-cell adhesion and biopolymeric extracellular matrix components to provide them with structure, as in the case of curli biofilm (Fig. 6a). Accordingly, we created PQN4 variants that produced various combinations of CsgF-IDP fusions, soluble secreted IDPs (“Sec-FUSLC”, “Sec-RGG”), and CsgA-IDP fusions (“Sec-FUSLC-CsgA”, “Sec-RGG-CsgA”) simultaneously (Fig. 6b-c). CsgA is the main structural component of native curli fibers, and assembles into amyloid fibers after secretion^40^. We envisioned that the soluble secreted IDPs could serve to bridge condensates forming at the cell surface and lead to larger aggregates. Similarly, we thought that CsgA-IDP fusions could serve as a mechanically robust anchor on which IDP condensates could form. We also created strains that produced either a CsgF fusion to (SIM)_5_ or a soluble secreted multivalent (SUMO)_5_ protein (Fig. 6d). (SUMO)_5_ and (SIM)_5_ are known to bind to each other in a multivalent manner^41^. This allowed us to explore a division of labor in producing cellular spatial patterns with one cell type engaging in IDP cell surface display (CsgF-(SIM)_5_) and another cell type producing a soluble multivalent molecular glue (SUMO)_5_. We verified the surface display of (SIM)_5_ and secretion of soluble (SUMO)_5_ using immunofluorescence staining images and flow cytometry (Extended Data Fig. 5a-c).

**Fig. 6.**
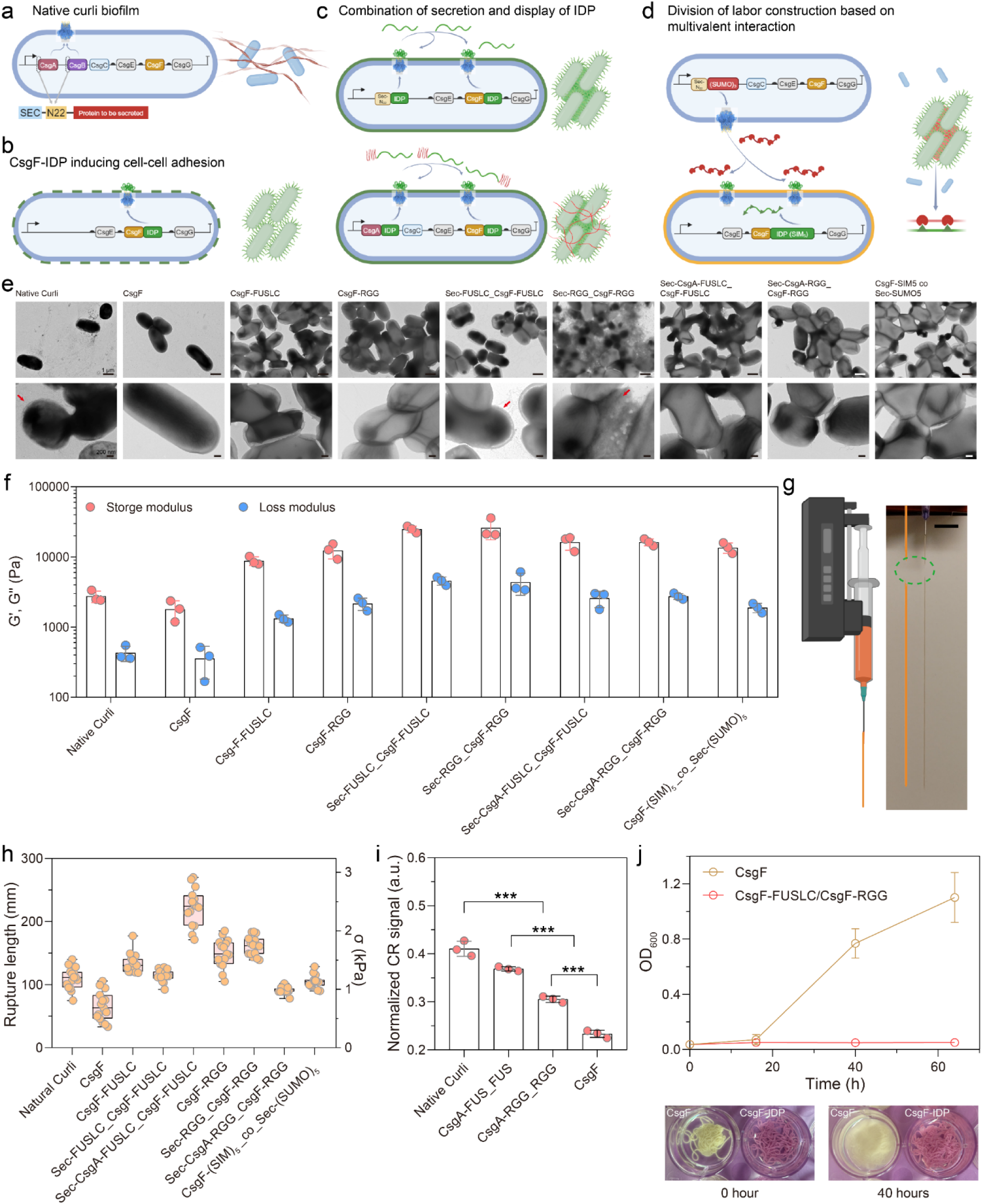
Combining IDP surface display with soluble IDP secretion or amyloid-scaffolded IDPs to build ELMs with tunable mechanical properties. **a-c**, Schematic depicting various engineered PQN4 strains, including CsgA only (i.e., native curli biofilm) (**a**), CsgF-IDP cell surface display (**b**), combined CsgF-IDP display and secreted soluble IDPs (**c**, top) or CsgA-IDP fusions (**c**, bottom). **d**, Schematic of system where one cell produces a non-self-aggregating CsgF-IDP fusion and another produces a multivalent “glue” scaffold in soluble form to cooperatively produce an ELM. **e**, Representative TEM images showing the morphologies of all ELMs. Red arrows refer point to the formation of extracellular protein aggregates. Scale bar: 1 μm (top) and 200 nm (bottom). **f**, Storage (G’) and loss (G”) moduli of all cell aggregate combinations at an angular frequency of 10 rad s^−1^ (n=3 repeats per group). **g**, Schematic depicting the syringe extrusion of an *E. coli* cell pellet to test for tensile properties (left). Photograph of setup, mid-experiment, (right), capturing the moment of filament rupture during extrusion. Scale bar: 2 cm. **h**, Rupture length and calculated tensile strength σ_exp_ of the extruded filament for cell pellets obtained from different PQN4 strain combinations (n=15 repeats per group). **i**, Congo Red pull-down assays quantifying amyloid produced for strains harboring plasmids encoding CsgA variants, compared to CsgF alone (i.e., no curli production) (n=3 repeats per group). **j**, Stability of ELM filaments immersed in PBS after extrusion. Stability was monitored by cellular shedding into the PBS solution using OD_600_ of the supernatant over time (n=3 repeats per group) for CsgF only and CsgF-IDP filaments. Images of filament mounds shown as pictures in the inset. P values were determined by one-way ANOVA with Tukey’s post-test; *P < 0.01, **P < 0.01, ***P < 0.001.

We started by using transmission electron microscopy (TEM) to investigate differences in microstructure for cellular aggregates that combine iDP^2^ with other scaffolding strategies. TEM images showed that cells displaying IDPs on their own were able to create large contact area cell-cell junctions (Fig. 6e). The combination of CsgF-IDP display and secreted soluble IDPs led to enhanced thickness for the cell-cell junctions, presumably with the soluble IDPs bridging the gap between cells spaced farther apart. The combination of CsgF-IDP and CsgA-IDP fusions also showed aggregates with more spaced-out cells and thicker ECM-like bridging structures. We studied the mechanical properties of these aggregates with conventional oscillatory shear rheology. Frequency sweep experiments (Extended Data Fig. 6a) showed that all the strain combinations we tried were viscoelastic solids throughout the tested range of angular frequency. Storage and loss modulus values were both 3-4 fold higher for CsgF-IDP compared with CsgA production alone (i.e., “native curli”) at a representative angular frequency of 10 rad/s (Fig. 6f). Co-secretion of CsgF-IDP fusions with CsgA-IDP fusions led to further increases in storage and loss modulus, similar to those observed for the CsgF-(SIM)_5_-(SUMO)_5_ combination. The highest modulus values were observed for the co-secretion of CsgF-IDPs with soluble IDPs.

We also characterized the tensile strength of the cellular aggregates using a syringe extrusion assay published previously^36^, where the stress near the extrusion nozzle increases as the hanging filament gets longer (Fig. 6g). Both displayed and secreted IDPs boost the tensile strength compared to strains without any IDPs and even with native curli alone. Interestingly, co-secretion of CsgA-FUSLC along with CsgF-FUSLC enhances tensile strength, but co-secretion of CsgA-RGG with CsgF-RGG showed a decrease in tensile strength compared to CsgF-RGG alone (Fig. 6h). This difference could be explained by lower amyloid formation measured for CsgA-RGG compared to CsgA-FUSLC according to a Congo Red assay (Fig.6i). Additionally, the segregation of immiscible cells types was preserved even after filament extrusion (Extended Data Fig. 6b-c), and the stability also showed durability compared CsgF only (Fig. 6j). Consequently, iDP^2^ is an empowerment to mediate the material properties of ELM.

## Discussion

Here we developed a new cell-surface display and cell patterning system for *E. coli*, called iDP^2^, based on the curli (Type VIII) secretion system and the fusion of IDPs to the surface anchor CsgF. iDP^2^ exhibited a very high density of cell-surface display, comparable with the highest values reported for other similar systems in the literature to date. Our approach is compatible with a wide range of unstructured protein domains but does not display structured domains as efficiently. The high surface density of the displayed domains promoted cellular aggregation during sedimentation from suspension, based on the propensity for certain IDP sequences to undergo liquid-liquid phase separation at high concentrations. A brief but systematic study of sequence dependency suggested the highest degree of compatibility with small, hydrophilic, or charged residues. Hydrophobic and bulky residues were less well tolerated and sequences with a high frequency of positively charged residues could not be displayed at all. Furthermore, the spatial patterning of cells within these aggregates could be programmed based on the distinct phase separation capabilities of the fused domains, with the CsgF-FUSLC and CsgF-RGG cells self-segregating to form island-like patterns of homogenous cell aggregates. The cell aggregation and patterning could be modulated by the known environmental responsiveness of the IDP domains, leading to cellular aggregates that exhibited temperature, and ion concentration-responsive behaviors over multiple cycles of modulation. Cell pellets held together by IDP-IDP interactions exhibited enhanced rheological properties and could be extruded to form self-standing filaments.

Overall, we see iDP^2^ as unique sub-class of CAMs, where the weak multivalent forces that drive IDP phase separation can be easily reconfigured based on environmental conditions compared to other CAMs based on specific molecular recognition (e.g., nanobody-antigen). The ability to form patterns directly from cell culture with minimal top-down processing make iDP^2^ a potentially useful tool for ELM fabrication that mimics certain aspects of structure building observed in nature. Our results also suggest that there may be a rich field of exploration for further leveraging IDP sequence to program the formation of other binary, tertiary, or even more complex patterns. Notably, cell surface display is a critical component of many high-throughput screening and directed evolution methods. So far, ELM research has been recalcitrant to high throughput techniques that have led to advances in other fields of biotechnology. Perhaps iDP^2^ could contribute to bridging this gap. The iDP^2^ strategy may also be broadly helpful to other synthetic biologists interested in using spatial organization to facilitate co-culture of different microbes in useful ways.

## Methods

### General reagent, instrument, and resources

Most chemical reagents were purchased from Sigma or Thermo Fisher unless stated otherwise. Kanamycin, Lysogeny broth (LB) granulated, and LB Agar were purchased from RPI Inc. All primers for PCR used in this work were ordered from Genewiz. Most plasmids containing protein to be used for surface display study were ordered from Addgene. Protein sequences, plasmids and strains were used in this work as shown in Table S4-6, respectively. General instruments include MiliQ, Nanodrop (Thermo Fisher), Centrigues (Eppendorf, Beckman), Oscillating incubator (INFORMS HT), Static incubator (Thermo Fisher).

### Construction of strains and plasmids

All plasmids containing some curli operon genes were constructed using the plasmid backbone from pBbB8k encoding CsgBACEFG prepared in our previous work. For the construction of iDP^2^ and evaluating the protein surface display, deletion of CsgBAC and integration of additional fusion sequence (including diverse tags, segments and whole proteins, as shown in Table S4) at the C-terminal of CsgF was conducted via PCR (with the Q5 polymerase New England Biolabs) (Applied Biosystems ProFlex PCR System) or double digest with standard restriction enzyme (New England Biolabs) to prepare desirable fragments, and further Gibson assembly (New England BIolabs, Gibson Assembly® Master Mix). Cloning was performed in the NEB® 5-alpha Competent *E. coli* strain selecting on plates with 50 µg/mL kanamycin (RPI). Plasmids were then harvested using the Qiagen Miniprep kit and sent for confirmatory sequencing with Genewiz or Plasmidsaurus Inc. Each plasmid variant was transformed into PQN4 cells, an *E. coli* cell strain derived from LSR10 (MC4100, ΔcsgA, λ(DE3), CamR) with the deletion of curli operon (ΔcsgBACEFG), using electroporation. Electrocompetent PQN4 cells were mixed with plasmid DNA before being subjected to a 1250-V pulse (by ECM 399 EXPONENTIAL DECAY WAVE ELECTROPORATION SYSTEM). The cells were immediately transferred to Super Optimal broth with Catabolite repression growth media (NEB) and were cultured for 1 h at 37 °C and 225 rpm before being plated on LB agar plates containing 50 µg/mL kanamycin.

### Cell Culture conditions

Cells were cultured in a shaking incubator (INFORS HT) at 37 °C and 225 rpm for the duration of the specified times. All cell cultures contained 50 μg/mL kanamycin to select for cells with the engineered plasmids. Single colonies taken from LB agar plates were used to make 5 mL of uninduced starter cultures that were grown overnight for all experiments to ensure stationary phase and consistent final density across samples. Cells were then diluted 100-fold into fresh media and grown for 4 hours before induction with L (+)-arabinose (final concentration of 0.1%, Thermo Scientific), and grown for additional 4 hours for most of study unless specifically noted in this work.

### Immunofluorescence staining for flow cytometry and confocal imaging

After culture and induction of all strains to be tested, samples were washed twice by dilution of 300 µL strain solution (normalized optical density (OD) 1) with 1 mL PBS in 1.5 mL tubes, centrifugation (5000 rpm for 4 min), and supernatant removal. After that, the cell pellets were suspended with 1-mL BD Pharmingen™ Stain Buffer (FBS) and centrifuged in the same condition. Final pellets were collected for the further immunofluorescence staining.

For Flow cytometry, each sample was treated with 100 µL APC anti-His Tag Antibody (BioLegend #362605, 1:40 dilution) for His tag staining, PE anti-FLAG (DYKDDDDK) Tag Antibody (BioLegend #637310, 1:40 dilution) for FLAG tag staining, or both together respectively for 30-minutes incubation at 4 °C. After staining, the samples were washed twice by mixing with 900 µL PBS for washing, then centrifugation to remove residual antibody in the supernatant. Finally, all groups of cell pellets were resuspended into 1 mL 1% Paraformaldehyde (PFA) fix solution (Epredia). The optional DAPI (Sigma) of 0.1 mg/mL (i.e., around 300 nM) staining was set up for 15 min before flow cytometry (Beckman Coulter CytoFLEX) test. For Spy tag characterization, the Spy Catcher (Biorad, # TZC002CYS3) labelled with FITC (Sigma) was used for covalent conjugation via Spy Tag-Spy Catcher isopeptide bond formation. Specifically, after washing 1 optical density unit of CsgF-Spy tag and control strains were incubated with excess Spy Catcher-FITC for 1 hour at 37 °C. Then, samples were washed with PBS three times, subjected to 1% PFA fixing, and treated with DAPI staining successively.

For confocal laser scanning microscopy (CLSM), the cell pellets were suspended onto polylysine pretreated glass slides (Histology Control System, INC.) for 5 min., and redundant suspension was discarded. Then, the attached cells were fixed by addition of 4% PFA for 2 hours, and washed with PBS two times. The slides with fixed cells were then blocked in 2% BSA (Sigma) solution for 1 hour at 37 °C. After washing, 200 µL Alexa488 anti-His Tag Antibody (Thermo Fisher, #MA1-21315-A488, 1:100 dilution) for His tag staining, Alexa594 anti-FLAG (DYKDDDDK) Tag Antibody (Novus Biologicals™ 1:100 dilution) for FLAG tag staining, or both was added for overnight incubation at 4 °C. After being washed 3 times with PBS, the slides were stained with 300 nM DAPI for 15 min, then were mounted on ProLong™ Diamond Antifade Mountant (Thermo Fisher), and covered with appropriate coverslips (Surgipath). Images were acquired using the Zeiss LSM 880.

### Fraction extraction of whole cell

Overnight induced PQN4 cells containing engineered plasmid variants were centrifuged at 4000 × g, and 4 °C, for 10 min after 24 h of growth. The 10-mL LB supernatant was taken to be the secreted medium fraction. To demonstrate the protein expression by western blot (as shown in Extended Data Fig. 1c), the cell pellets were lysed by lysis buffer from BugBuster@ Master Mix (Sigma), containing 0.01 mg/mL Benzonase (Sigma) and 0.1% protease inhibitor (Thermo Fisher), for 30 minutes on ice. The lysis solution was centrifuged at 15,000 g for 10 minutes at 4 °C. The supernatant was collected as cellular fraction.

For distinguishing the cellular fractions (as shown in Extended Data Fig. 2a-c), after separating cell pellet and secreted medium, a periplasmic extraction was performed using a 50 mM Tris/0.53 µM EDTA/20% Sucrose (TES) buffer based on a protocol optimized by Ghamghami et al^42^. Cell pellets were resuspended in 0.5 mLof cold TES buffer and were incubated on ice for 1 h. The suspensions were spun down at 15 000 × g at 4 °C for 30 min, and the supernatant was taken to be the periplasmic protein fraction. The pellets were resuspended in 1mL lysis buffer of BugBuster@ Master Mix (Sigma) containing 0.01 mg/mL Benzonase (Sigma) and 0.1% protease inhibitor (Thermo Fisher). The samples were incubated on ice for 20 minutes then centrifuged at 15,000 x g for 10 minutes at 4 °C. The supernatant containing soluble cytosolic proteins was collected for analysis. The pellets were resuspended in 1 mL 2% Triton X-100 (SPECTRUM CHEM) to solubilize membranes and membrane proteins, followed by centrifugation at 8,000 x g for 20 minutes at 4 °C. The supernatant containing membranes and membrane proteins was saved as membrane fraction. The remaining pellets were resuspended in 8 M urea to solubilize insoluble proteins in inclusion bodies. The samples were centrifuged at 8,000 x g for 20 minutes at 4 °C and supernatant containing solubilized inclusion body proteins was collected for analysis. For downstream applications (i.e., SDS-PAGE, Western Blot, etc.), each fraction was diluted back to the original culture volume is necessary. For example, for a 10 mL culture: Secreted fraction was not diluted, Periplasmic fraction was diluted 20X, Cytosolic fraction was diluted 10X, Inclusion body fraction was diluted 10X.

### CsgF-His tag purification

According to published literature^27^, recombinant CsgF with C-terminal His-tag (Sequence as shown in Table S4) of pET21d backbone was overexpressed in *E. coli* BL21(DE3) (New England Biolabs) using 500 µM IPTG for 4 h at 37 °C. The cell pellet from 250 mL culture was lysed in 25 mL of 8 M GdmCl, 150 mM NaCl, and 50 mM potassium phosphate (KPi) pH 7.3, and the denatured solution was kept on a shaker overnight at room temperature (RT). The lysate was centrifuged, and the supernatant was mixed with 3 mL of Ni-NTA beads. These were allowed to bind for an hour at RT, and further purified via HisPur™ Ni-NTA Spin Columns (Thermo Fisher). Ni-NTA beads were washed with 9 different solutions as below: Wash 1: 10 mL of 8 M urea, 650 mM NaCl, 50 mM KPi pH 7.3. Wash 2: 10 mL of 2 M urea, 650 mM NaCl, 50 mM KPi pH 7.3. Wash 3: 10 mL of 2 M urea, 20 mM imidazole, 650 mM NaCl, 50 mM KPi pH 7.3. Wash 4: 10 mL of 2 M urea, 40 mM imidazole, 650 mM NaCl, 50 mM KPi pH 7.3. Wash 5: 10 mL of 2 M urea, 60 mM imidazole, 650 mM NaCl, 50 mM KPi pH 7.3. Wash 6: 10 mL of 2 M urea, 80 mM imidazole, 650 mM NaCl, 50 mM KPi pH 7.3. Wash 7: 10 mL of 2 M urea, 100 mM imidazole, 650 mM NaCl, 50 mM KPi pH 7.3. Wash 8: 10 mL of 2 M urea, 200 mM imidazole, 650 mM NaCl, 50 mM KPi pH 7.3. Wash 9: 10 mL of 2 M urea, 500 mM imidazole, 650 mM NaCl, 50 mM KPi pH 7.3.

The eluted solutions were collected respectively for SDS-PAGE, which is used for the confirmation of purified CsgF-His tag. As shown in Extended Data Fig. 3d, the eluents of wash 7-9 containing purified proteins were dialyzed against 1% acetic acid and lyophilized (LABCONCO). The protein was stored at −20 °C for further use.

### Western-blot and quantification

Each fractioned sample and purified CsgF sample was loaded onto the 4–20% Mini-PROTEAN® TGX™ Precast Protein Gels (Bio-Rad) with specified concentration. After running, the gels were transferred to polyvinylidene difluoride membranes using the iBlot 2 transfer device (Invitrogen) and corresponding protocol. Then, membranes were washed with 1×Tris-buffered saline/Tween (TBST, Thermo Fisher) solution and placed in a 5% BSA (Sigma) blocking solution for 1 h. Furthermore, the blocked membranes were incubated with anti-His tag antibody with HRP conjugate (Invitrogen) in 1% BSA at 4 °C overnight with gentle rocking. After washing 3 times in 1x TBST for 10 minutes each, the membranes were transferred to a solution containing the enhanced chemiluminescence substrate (Pierce™ ECL Western Blotting Substrate, Thermo Fisher) to allow visualization of His-tagged proteins. The blotted proteins were visualized in a ChemiDoc Imaging System (Bio-Rad) using a chemiluminescence detector setting. The quantitative western blot was based on the band quantification of integral of pixel value by Image J. The purified CsgF samples with specified concentrations were selected as standard samples for standard curve fitting according to selected linear region.

### CFU test

Induced strains were normalized to OD_600_ 1, serving as the initial culture. The initial culture was diluted into fresh LB 1:10 (100 µL culture + 900 µL LB). 1:10 serial dilutions were completed into 7 tubes with triplicates. 10 µL of the cells was added dropwise onto the agar plates containing 50 µg/mL kanamycin and incubate overnight. Visualization of colonies growth on the plates was acquired by ChemiDoc Imaging System (Bio-Rad) as shown in Extended Data Fig. 2e-f. The number of colonies was counted from different dilutions to calculate their cellular concentration and ensure consistent results.

### Standard particles for the quantified flow cytometry

Commercial Ultra Rainbow & Supra Rainbow Quantitative Particle Kits (Spherotech) were used for quantitative flow cytometry. The standard particles with different fluorescent intensity have been assigned ERF (Equivalent Number of Reference Fluorophores) values. ERF versus MFI of APC could be utilized for the standard curve. The same flow conditions were used for running the cellular samples stained with APC-anti His tag antibody and further quantification.

### Aggregation and sedimentation assay

Induced strains as mentioned before were collected and suspended into PBS after washing twice with PBS. 3-mL cell solution for each group were transferred into the 5 mL sterile glass tubes, then kept at different temperatures (4 °C, 25 °C, 37 °C) overnight. After incubation, samples of 150 µL were taken from the top 25% of the culture solution (supernatant) into 96-well assay plates and OD_600_ was measured on the SpectraMax microplate reader (Molecular Devices), with triplicates for each group. Additionally, visible pictures were acquired of these samples. The PBS was also replaced with buffer containing different concentrations of NaCl (0, 50, 100, 150, 300 mM, pH 7.4) to test the variation of aggregation intensity of different strains. The normalized aggregation intensity was calculated via Normalized aggregation intensity (%) = 100 x 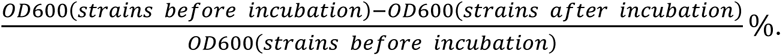

### Ionic responsibility of aggregation

Following the aggregation study, induced strains were repeatedly suspended or switched into different culture medium of 0 mM NaCl and 50 mM NaCl. After each culture medium change, samples were incubated at 4 °C for 12 hours. The samples for OD_600_ measurement were acquired by the same method as mentioned in aggregation assay.

### Microscopy study for the pattern formation

The strains with intracellular fluorescent protein expression labelling were prepared by the transformation of plasmids with both fluorescent protein gene and optimized curli operon. These were cultured following the normal condition and induction procedure. Several strains were prepared including CsgF (CFP/RFP), CsgF-FUSLC (CFP/YFP/RFP), CsgF-RGG (CFP/RFP), CsgF-FUSLC_CsgF-RGG (YFP), CsgF-ELP(S48I48) (CFP), CsgF-ELP(S96) (CFP). All strains were normalized to OD_600_ 1 before preparing 2-component (1:1) and 3-component (1:1:1) mixtures. After mixing based on the different designed combination of strains, the mixtures in the centrifuge tubes were vortexed briefly to ensure a homogeneous solution. 400ul of the mixtures were transferred into the 8-well plates (Thermo Scientific™ Nunc™ Lab-Tek™ Chambered Coverglass) and further incubated at 4 °C overnight. The plates were directly used for imaging, including the 2D snapshots and 3D Z-stack.

### Analysis of pattern formation via BiofilmQ

BiofilmQ, an open-source software written in MALTALB (R2022b), was used for biofilm analysis of 3D images from CLSM. The biofilm segmentation area is dissected into small squares (with length of 1.2 μm), analogous to the cubes in 3D images. After segmentation, diverse biofilm parameter quantification and data visualization, including nearest neighbor distance, local density, local thickness, and biofilm toughness, were calculated via the algorithms described in detail with examples in the documentation provided online (https://drescherlab.org/data/biofilmQ).

### Nearest neighbor distance ratio definition and analysis

The nearest neighbor distance ratio (*r*_i_) was defined as the ratio of the closest in-phase neighbor and the closest out-of-phase distance neighbor, to characterize the mixed distribution uniformity of two-phase strains as mentioned. The designed formula of (*r*_i_) was incorporated into BiofilmQ and ran for the data analysis.

### Dynamic pattern formation study

Similar to the study of ionic responsibility aggregation, the mixture of CsgF-FUSLC (CFP) and CsgF-RGG (RFP) were used for repeated suspension and switching between two different culture mediums of 0 mM NaCl and 50 mM NaCl. At each time point, samples were incubated in the 8-well plates overnight and collected for further imaging study. The 3D images were collected and further BiofilmQ analysis performed on the samples to achieve visualization and quantification of dynamic pattern formation as well as the nearest neighbor distance ratio, respectively.

### Cellular condensation-based ELM preparation

1 L cultures of PQN4 strains transformed with various iDP^2^-based plasmids were grown and induced overnight to ensure sufficient protein expression. Cellular pellet of ELM was collected by subjecting the culture to filtration using a Mixed Cellulose filter membrane with 0.22 um GSWP (Sigma) or centrifugation at 5000 rpm for 20 minutes. The collected ELM were then washed once with 10 mL PBS supplemented with Benzonase, and then three times with 10 mL Milli-Q water. The collected ELM was then transferred into tubes for further study.

### Transmission electron microscopy (TEM) for the cellular condensation morphology

The collected cellular pellets were suspended in the Milli-Q water and dropped onto carbon-coated copper grids (Electron Microscopy Sciences). The excess liquid on the copper grids was blotted off with filter paper (Cytiva). Then, the grids were allowed to air dry and examined on a TEM (JEOL JEM 1010).

### Rheology study

The ELM samples were loaded on the rheometer using a plate and plate system (40 mm) of Discovery Hybrid Rheometer-3 and TRIOS software (TA Instruments, New Castle, DE). Strain sweeps were carried out with shear strain from 0.01% to 100% at a frequency of 10 rad/s. Frequency sweep experiments from 100 to 0.01 rad/s were performed at a 0.2% strain amplitude. Data obtained from at least three independent samples was reported.

### Tensile strength study

The ELM samples were transferred into a 1-mL syringe. Syringe tips were sealed, and the cell pellet was spun down at 5,000 × g for 5 min. to remove bubbles. A syringe pump (with flow of 0.1 mL/min) was used for the extrusion of materials from the syringe with a blunt syringe needle of 21 gauge (BENECREAT). The length of extruded materials was measured by the pre-configured rulers and recorded when material string broke. According to the previous literature^36^, the ruptured length was used to determine the tensile length 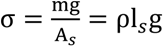, with the gravitational force mg, g = 9.81 m/s^2^, the cross-sectional area A_s_, the length l_s_, and the volume V_s_ = *A*_s_l_s_ of extruded material string below the breaking point.

### Congo Red Pull Down Assay

1 mL induced strains were spun down at 6000 rpm for 10 minutes. The supernatant was discarded and sample resuspended in 1 mL 0.05% Congo red (Sigma) solution. After 10-minutes incubation, the samples were centrifuged for 10 minutes at 14,000 rpm. The supernatant was collected for absorbance measurement at 490 nm. Also, the original culture cells were collected for OD_600_ measurement. The signal was subtracted from a Congo Red blank, divided by the culture’s OD_600_ measurement to reflect curli production per cell.

### Macroscopic ELM study

The 1 mL extruded material string was put into the well plates and immersed into the PBS for different strains. The static incubation at RT was set up and the supernatant was collected the at different time points to evaluate the stability of ELM via OD_600_ measurement. The material mixture of CsgF-FUSLC (CFP) and CsgF-RGG (RFP) was used for the images captured by microscopy (Revolve).

## Acknowledgments

We really appreciate the Institute for Chemical Imaging of Living Systems (CILS) of Northeastern University and Dr. Guo-Xin Rong’s help with Confocal and Flow cytometry study. We really appreciate the Boston Electron Microscopy Center (BEMC) of Northeastern University, William Fowle, and Dr. Shirin Kaboli for their assistance of TEM study. We thank Michelle Issac for her support with PCR and plasmids preparation. Parts of the schematics were adapted from BioRender.com (https://BioRender.com/y65u777).

## Author contributions

Conceptualization: R.C. and N.S.J. Methodology: R.C., H.T. and N.S.J. Investigation: R.C. and H.T. Visualization: R.C., H.T. and N.S.J. Funding acquisition: N.S.J. Project administration: R.C. Supervision: R.C. and N.S.J. Writing – original draft: R.C. Writing – review & editing: R.C., H.T. and N.S.J.

## Data and materials availability

All data are available in the main text or the supplementary materials.

## Supplementary Information

Supplementary Text (Note S1) References

---

**Extended Data Fig. 1.**
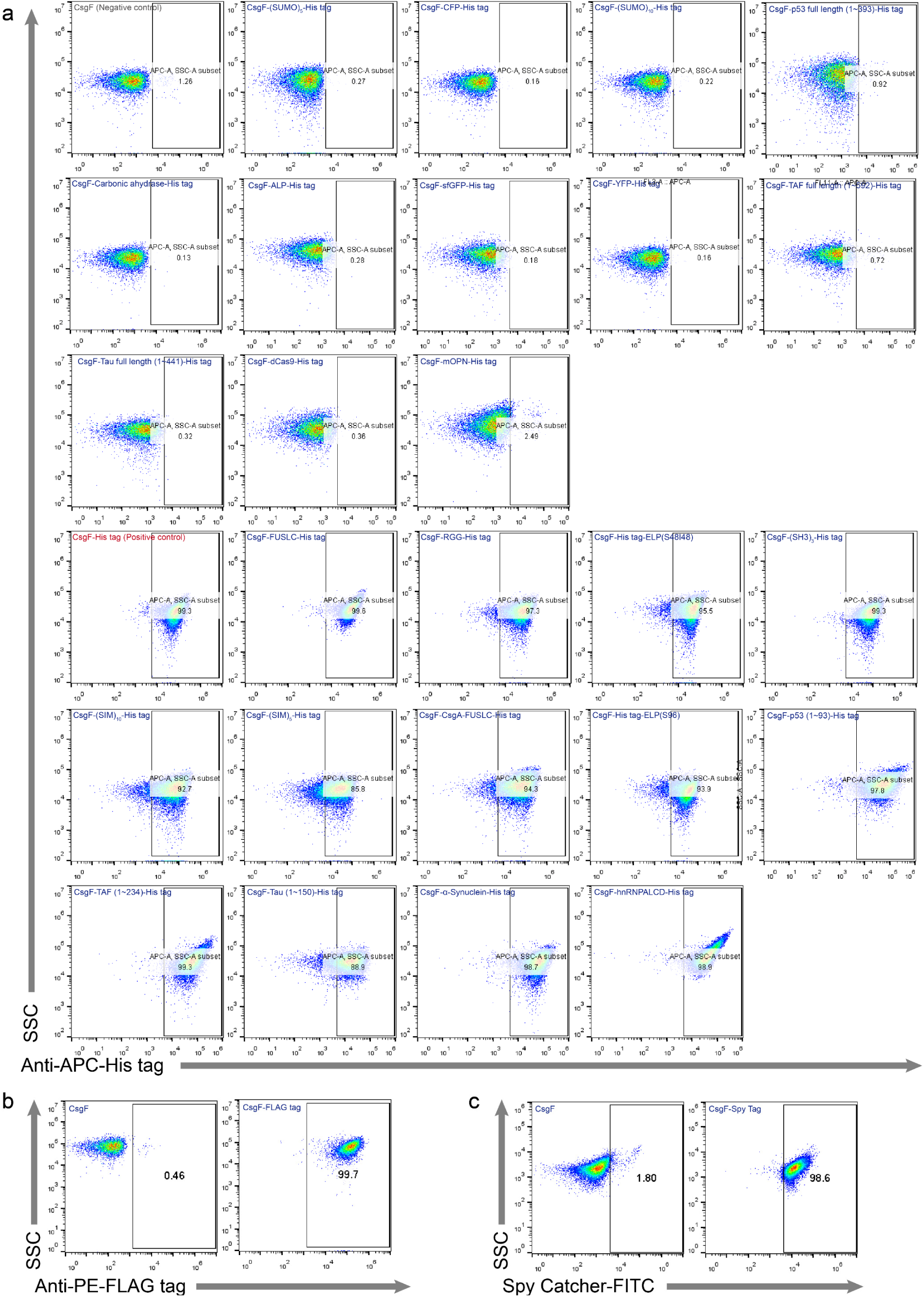
**a**, Representative anti-His tag-APC versus SSC-A dot plots for PQN4 displaying CsgF fused with protein of interest (POI), gated region based on negative control of CsgF only strain and positive control of CsgF-His tag strain. **b**, Representative anti-FLAG tag-PE versus SSC-A dot plots for PQN4 displaying FLAG tag or not. **c**, Representative Spy Catcher-FITC versus SSC-A dot plots for PQN4 displaying Spy tag or not.

**Extended Data Fig. 2.**
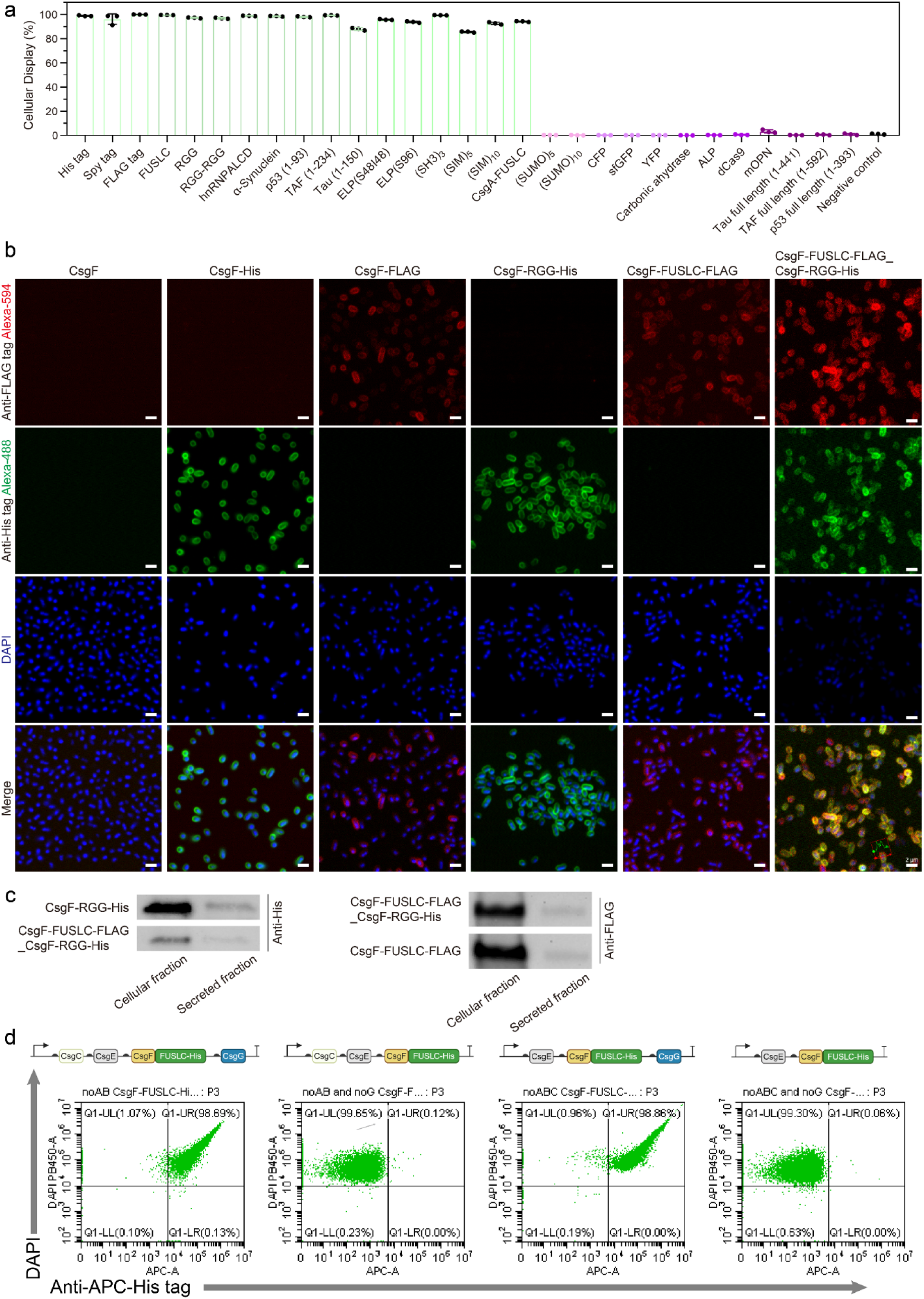
**a**, Percent of cellular display for the protein sequences library including diverse folded sequence and IDP. **b-c**, CLSM images (**b**), western blot (**c**) of PQN4 displaying representative IDPs, including FUSLC, RGG or both FUSLC and RGG together, His tag and FLAG tag respectively fused with FUSLC and RGG for immunofluorescence staining, only CsgF as negative control. Scale bars of (**b**): 2 μm. **d**, Scatter plot of flow cytometry for different constructs with CsgC, CsgG, both, or neither.

**Extended Data Fig. 3.**
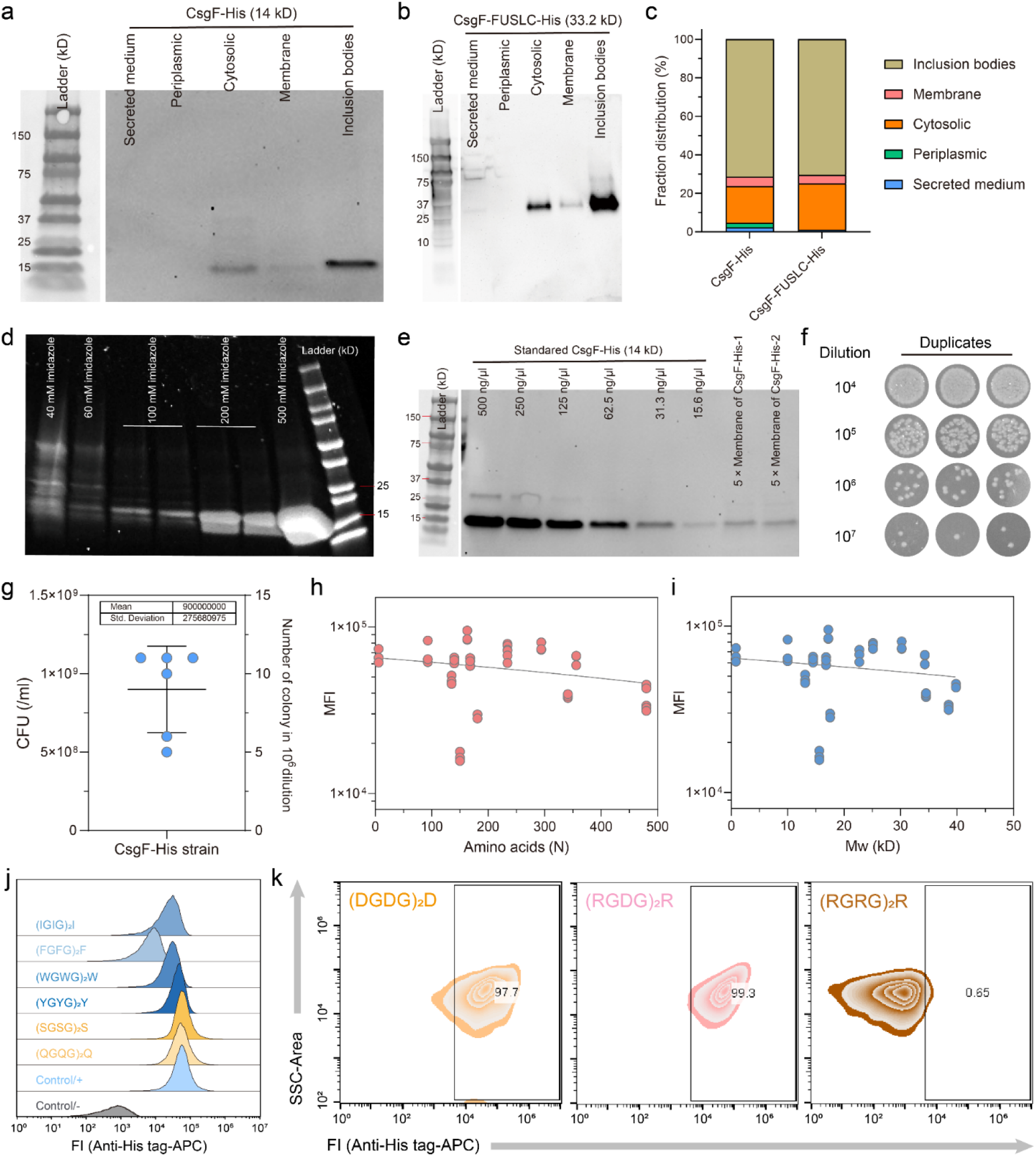
**a-c**, Western blot of cellular fraction for the strains of CsgF-His tag (**a**) and CsgF-FUSLC-His tag (**b**), and their quantitative percentage analysis (**c**). **d**, Coomassie blue staining SDS-PAGE gel of CsgF-His tag purification by elution with gradient concentration of imidazole solution. **e**, Western blot of purified CsgF-His tag with different concentrations and cellular membrane fraction. **f-g**, CFU assay to quantify the cellular concentration by dilution (**f**) and their statistical results (**g**). **h-i**, MFI of flow cytometry for diverse display of IDPs with different amino acids number (**h**) and molecular weight (**i**). **j**, Histogram of fluorescent intensity of different strains with designed sequences display. **k**, Representative anti-His tag-APC versus SSC-A dot plots for PQN4 displaying CsgF fused with (DGDG)_2_R, (RGDG)_2_R, and (RGRG)_2_R.

**Extended Data Fig. 4.**
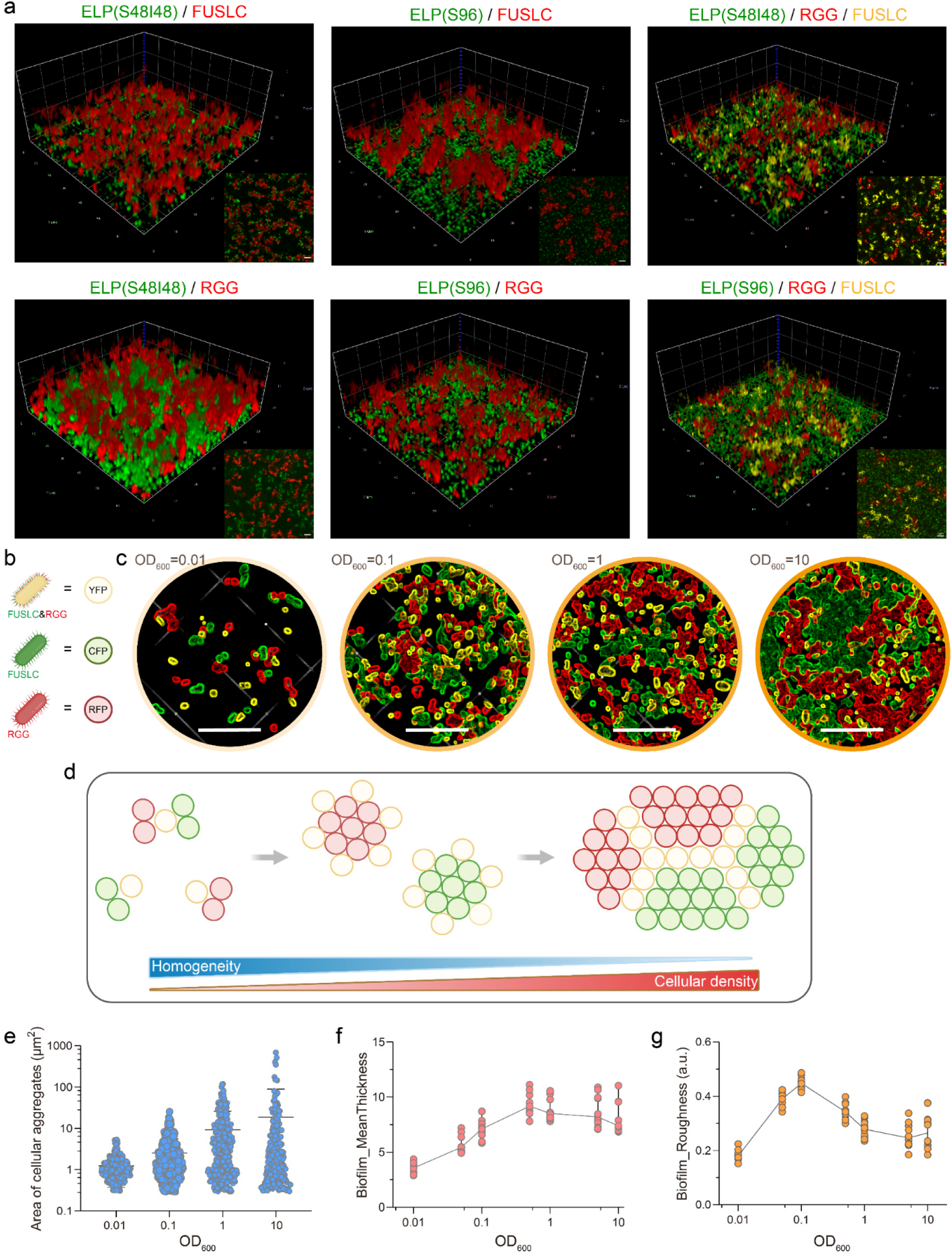
**a**, Various pattern formation based on strains (labeled with different fluorescent protein expression intracellularly) displaying different IDPs, including RGG, FUSLC and ELPs. **b**, Schematic strains with different IDPs display including FUSL, RGG, and both FUSLC and RGG together. **c**, cellular distribution and pattern formation in different cellular concentrations (OD_600_) for a three-strain system. Scale bar: 10 μm. **d**, Schematic pattern formation mediated by increase in cellular concentration. **e**, Area of cellular aggregation for the different strains’ concentrations via images analysis. (More than 200 data points from 3∼4 repeats of images). **f-g**, Biofilm thickness (**f**) and roughness (**g**) for the different strains’ concentrations via Biofilm Q. (n=9∼12 repeats per group).

**Extended Data Fig. 5.**
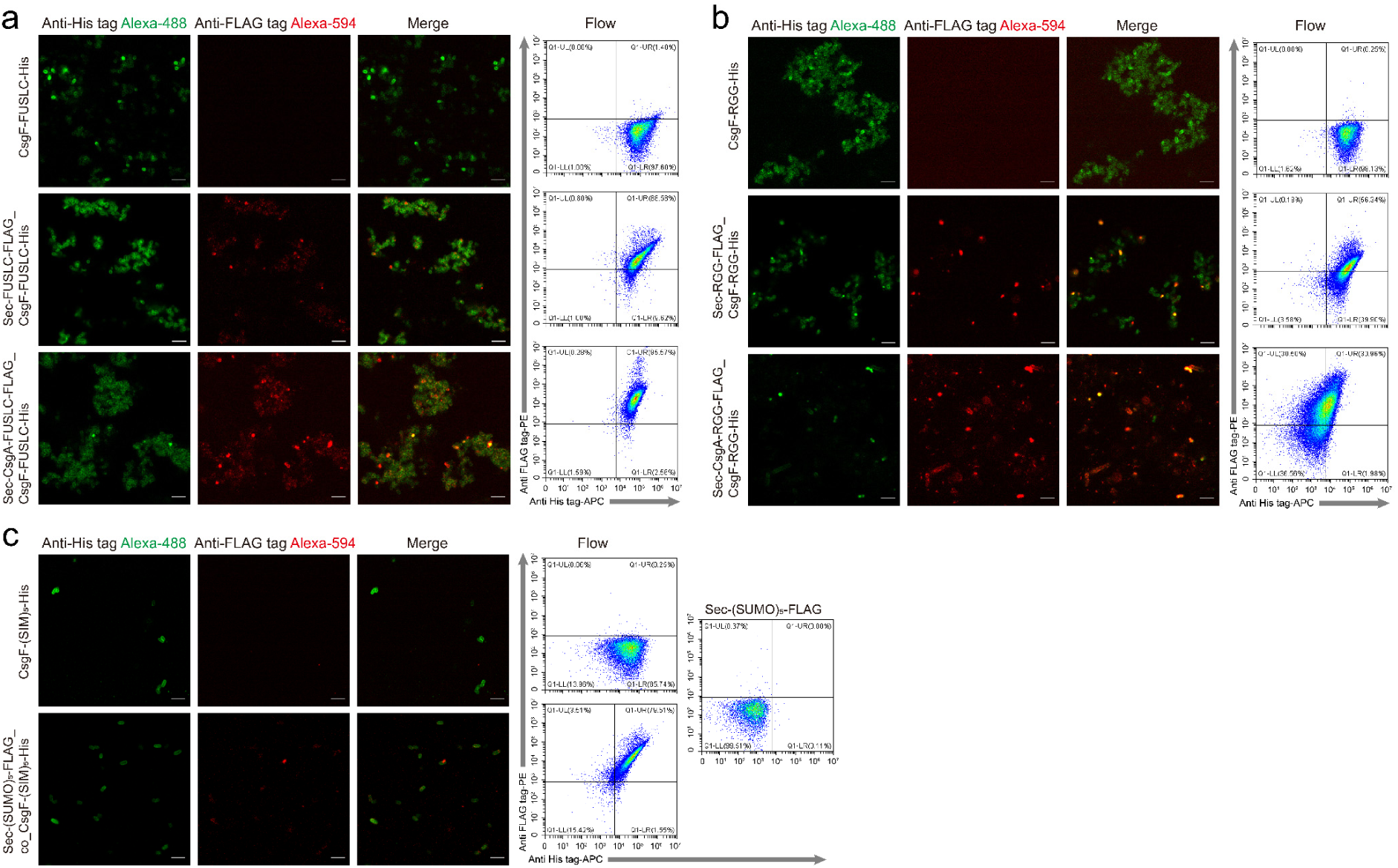
Immunofluorescence of CLSM and flow cytometry to prove the protein surface display and secretion to binding with membrane, including (**a**) CsgF-FUSLC, Sec-FUSLC_CsgF-FUSLC and Sec-FUSLC-CsgA_CsgF-FUSLC, (**b**) CsgF-RGG, Sec-RGG_CsgF-RGG and Sec-RGG-CsgA_CsgF-RGG, (**c**) CsgF-(SIM)_5_, Sec-(SUMO)_5__co_CsgF-(SIM)_5_ and flow of Sec-(SUMO)_5_. Scale bar of images: 2 μm.

**Extended Data Fig. 6.**
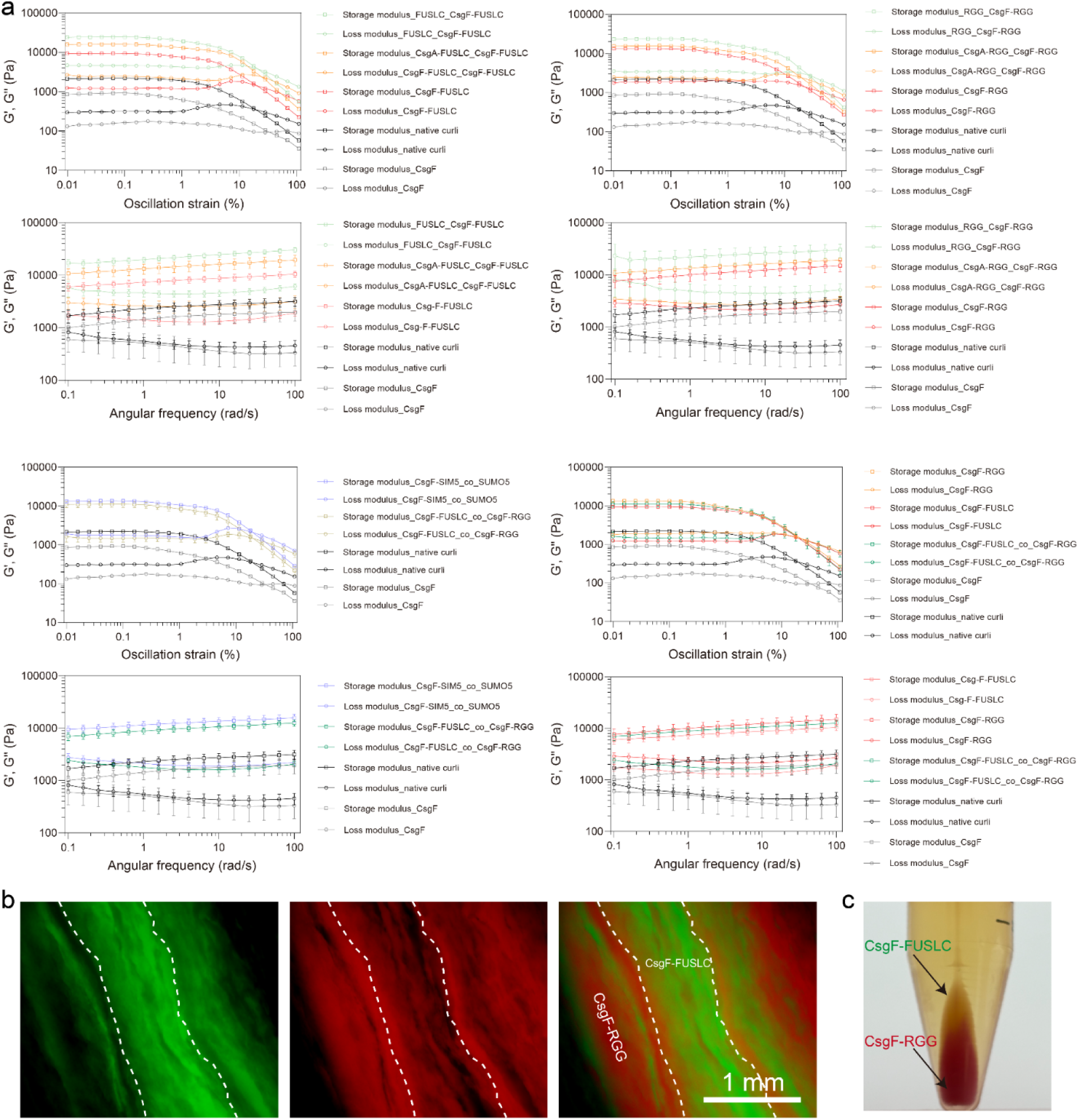
**a**, Oscillatory strain sweep and frequency sweep curve. Strain sweep measurements were acquired from 0.01% to 100% strain amplitude at a constant frequency of 10 rad/s. (n=3 repeats each group) From the amplitude sweep curves, we identified the linear viscoelastic region of the cellular pellets and set the strain used to collect frequency sweep data to 0.2%. Frequency sweep measurements were acquired from 0.1 rad/s to 100 rad/s at a constant strain amplitude of 0.2%. **b**, Images of macroscopic patterns with chaotic structure after extrusion from the syringe for the CsgF-FULLC/CsgF-RGG mixture. Scale bar: 1 mm. **c**, Pictures of CsgF-FULLC (green)/CsgF-RGG (red) mixture with clear interfacial separation after centrifuge.

