## Supplementary Text (Note S1) Supplementary Table (S1-S6) for "Programming cellular condensates for living materials using an intrinsically disordered protein display platform (iDP^2^)"

Note S1. Quantitative surface density of protein displays by western blot or flow cytometry of standard fluorescent particles

The standard curves of western blot of purified CsgF-His with different concentrations were fitted linearly, which provide the quantification of unknown CsgF-His samples from fractionation. The concentration of membrane fraction was 3.34±0.01 ng/uL from 10 optical density units. CFU assay was used for the measurement of cellular concentration of culture strains (9 x $10^8$ $mL^{-1}$) for fractionation. Therefore, the number of CsgF-His tag for each cell is (16.1 ± 4.9) × $10^4$ via sequential calculation of formula (1-2).

$$\text{Quantiy of surface protein for each cell (mg)} = \frac{Protein\ concentraion\ of\ membrane\ fraction\ (ng{\cdot}\mu l^{-1})}{1000{\cdot}CFU\ ml^{-1}} \cdots\cdots\cdots (1).$$

$$\text{Number of CsgF for each cell} = \frac{Quantiy\ of\ surface\ protein\ for\ each\ cell\ (mg) \cdot N_A}{1000 \cdot M_w} \cdots\cdots\cdots (2)$$

The standard curves of flow cytometry for standard particles with different equivalent number of reference fluorophores (ERF) values are shown in Fig. 3C-D The amount of CsgF-His per cell was calculated as (7.4 ± 0.4) ×$10^4$ via the standard curve.

Both western blot and flow cytometry provide the measurement of the number of CsgF-His tag each cell. Assuming that all cells are the same size and surface area, further surface density of displayed proteins could be computed using the surface area of cells via formula (3).

$$\text{CsgF density on the surface} = \frac{Number\ of\ CsgF\ for\ each\ cell}{A_{cell}} \cdots\cdots\cdots (3)$$

In theory, each curli secretion channel based on CsgE-CsgG-CsgF complex contains 9 CsgG and 9 CsgF. Accordingly, the CsgEGF complex density on the surface could also be computed via formula (4).

$$\text{CsgEGF complex density on the surface} = \frac{CsgF\ density\ on\ the\ surface}{9} \cdots\cdots\cdots (4)$$

Assuming that all curli secretion channels have completely homogeneous distribution on the extracellular membrane, the distance between 2 CsgEGF complex could also be computed via formula (6).

$$\text{Distanace between two CsgEGF complex} = \left(\frac{2 \cdot 100000}{\sqrt{3} \cdot CsgEGF\ complex\ density\ on\ the\ surface}\right)^{\frac{1}{2}} \cdots\cdots (6)$$

All calculated results are summarized in the Table S2-3.

**Table S1. Related constant for the calculations**

| | Description | Unite | Value |
|---|---|---|---|
| $N_A$ | Avogadro constant | $mol^{-1}$ | $6.02\times10^{23}$ |
| $M_w$ | Molecular weight of CsgF-His tag | g/mol (Da) | 13876 |
| $A_{cell}$ | Surface area of the cell | $\mu m^2$ | 4.5[1] |

**Table S2. Measured parameters of iDP$^2$ based on quantified western blot**

| Description | Mean ± SD |
|---|---|
| CsgF concentration of membrane (ng/μL) | 3.34±0.01 |
| CFU (/mL) | $(9.0\pm2.76)\times10^8$ |
| Quantity of CsgF for each cell (mg) | $(3.71\pm1.14)\times10^{-12}$ |
| Number of CsgF for each cell | $(16.1\pm4.9)\times10^4$ |
| CsgF density (/$\mu m^2$) | $(35.7\pm11.0)\times10^3$ |
| CsgEGF density (/$\mu m^2$) | $(3.97\pm1.22)\times10^3$ |
| Area of 3 CsgEFG complex ($nm^2$) | 755±231 |
| Distance between 2 CsgEFG (nm) | 17.0±2.6 |

**Table S3. Measured parameters of iDP$^2$ based on quantified Flow cytometry of standard particles**

| Description | Mean ± SD | | |
|---|---|---|---|
| | CsgF-His tag | CsgF-FUSLC-His tag | CsgF-RGG-His tag |
| Number of CsgF for each cell | $(7.4\pm0.4)\times10^4$ | $(8.1\pm0.2)\times10^4$ | $(6.2\pm0.2)\times10^4$ |
| CsgF density (/$\mu m^2$) | $(16.3\pm0.9)\times10^3$ | $(18.1\pm0.4)\times10^3$ | $(13.9\pm0.4)\times10^3$ |
| CsgEGF density (/$\mu m^2$) | $(1.82\pm0.10)\times10^3$ | $(2.01\pm0.04)\times10^3$ | $(1.54\pm0.05)\times10^3$ |
| Area of 3 CsgEFG complex ($nm^2$) | 1652±90 | 1494±31 | 1947±60 |
| Distance between 2 CsgEFG (nm) | 25.2±0.7 | 24.0±0.3 | 27.4±0.4 |

**Table S4. Protein sequence used in this work**

| Name | Sequence | Source and state |
|---|---|---|
| All curli operon related protein sequences | CsgA:<br>MKLLKVAAIAAIVFSGSALAGVVPQYGGGGNHGGGGNNSG<br>PNSELNIYQ<br>YGGGNSALALQTDARNSDLTITQHGGGNGADVGQGSDDSSI<br>DLTQRGFGNSA<br>TLDQWNGKNSEMTVKQFGGGNGAAVDQTASNSSVNVTQV<br>GFGNNATAHQY<br>***(Red: Sec, Green: N22)*** | Plasmid of Ref.[2] that lab has. |

| | | |
|---|---|---|
| | CsgB:<br>MKNKLLFMMLTILGAPGIAAAAGYDLANSEYNFAVNELSKS<br>SFNQAAIIG<br>QAGTNNSAQLRQGGSKLLAVVAQEGSSNRAKIDQTGDYNL<br>AYIDQAGSANDA<br>SISQGAYGNTAMIIQKGSGNKANITQYGTQKTAIVVQRQSQM<br>AIRVTQR<br>CsgC:<br>MNTLLLLAALSSQITFNTTQQGDVYTIIPEVTLTQSCLCRVQIL<br>SLREGSS<br>GQSQTKQEKTLSLPANQPIALTKLSLNISPDDRVKIVVTVSDG<br>QSLHLSQQWPP<br>SSEKS<br>CsgE:MKRYLRWIVAAEFLFAAGNLHAVEVEVPGLLTDHTVS<br>SIGHDFYRAFSD<br>KWESDYTGNLTINERPSARWGSWITITVNQDVIFQTFLFPLKR<br>DFEKTVVFALI<br>QTEEALNRRQINQALLSTGDLAHDEF<br>CsgF:<br>MRVKHAVVLLMLISPLSWAGTMTFQFRNPNFGGNPNNGAFL<br>LNSAQA<br>QNSYKDPSYNDDFGIETPSALDNFTQAIQSQILGGLLSNINTG<br>KPGRMVTNDYI<br>VDIANRDGQLQLNVTDRKTGQTSTIQVSGLQNNSTDF<br>CsgG:<br>MQRLFLLVAVMLLSGCLTAPPKEAARPTLMPRAQSYKDLTH<br>LPAPTGK<br>IFVSVYNIQDETGQFKPYPASNFSTAVPQSATAMLVTALKDS<br>RWFIPLERQGL<br>QNLLNERKIIRAAQENGTVAINNRIPLQSLTAANIMVEGSIIGY<br>ESNVKSGGVG<br>ARYFGIGADTQYQLDQIAVNLRVVNVSTGEILSSVNTSKTILS<br>YEVQAGVFRFID<br>YQRLLEGEVGYTSNEPVMLCLMSAIETGVIFLINDGIDRGLW<br>DLQNKAERQND<br>ILVKYRHMSVPPES | |
| CsgF-His tag | MGTMTFQFRNPNFGGNPNNGAFLLNSAQAQNSYKDPSYND<br>DFGIETPSALD<br>NFTQAIQSQILGGLLSNINTGKPGRMVTNDYIVDIANRDGQL<br>QLNVTDRKTGQ<br>TSTIQVSGLQNNSTDFGS**HHHHHH** | Plasmid of Ref.[2] that lab has. Sequence |

| | | |
|---|---|---|
| | *(Underline: His tag)* | reference ref.[3] |
| FALG Tag | DYKDDDDK | Extension primers of PCR |
| Spy tag | AHIVMVDAYKPTK | Extension primers of PCR |
| p53 | MEEPQSDPSVEPPLSQETFSDLWKLLPENNVLSPLPSQAMDD LMLSPDDIEQW FTEDPGPDEAPRMPEAAPPVAPAPAAPTPAAPAPAPSWPLSSS VPSQKTYQGS YGFRLGFLHSGTAKSVTCTYSPALNKMFCQLAKTCPVQLWV DSTPPPGTRVRA MAIYKQSQHMTEVVRRCPHHERCSDSDGLAPPQHLIRVEGN LRVEYLDDRNTF RHSVVVPYEPPEVGSDCTTIHYNYMCNSSCMGGMNRRPILTI ITLEDSSGNLLGR NSFEVRVCACPGRDRRTEEENLRKKGEPHHELPPGSTKRALP NNTSSSPQPKKKP LDGEYFTLQIRGRERFEMFRELNEALELKDAQAGKEPGGSRA HSSHLKSKKGQST SRHKKLMFKTEGPDSD *(Red: p53(1-93), black: p53(94-279), green: p53(280-393))* | Addgene Plasmid #24859 |
| TAF | MSDSGSYGQSGGEQQSYSTYGNPGSQGYGQASQSYSGYGQT TDSSYGQNYSG YSSYGQSQSGYSQSYGGYENQKQSSYSQQPYNNQGQQQNM ESSGSQGGRAP SYDQPDYGQQDSYDQQSGYDQHQGSYDEQSNYDQQHDSYS QNQQSYHSQR ENYSHHTQDDRRDVSRYGEDNRGYGGSQGGGRGRGGYDK DGRGPMTGSSG GDRGGFKNFGGHRDYGPRTDADSESDNSDNNTIFVQGLGEG VSTDQVGEFFK QIGIIKTNKKTGKPMINLYTDKDTGKPKGEATVSFDDPPSAK AAIDWFDGKEFH GNIIKVSFATRRPEFMRGGGSGGGRRGRGGYRGRGGFQGRG GDPKSGDWVC PNPSCGNMNFARRNSCNQCNEPRPEDSRPSGGDFRGRGYGG ERGYRGRGGR | Addgene Plasmid #84895 |

| | | |
|---|---|---|
| | GGDRGGYGGDRSGGGYGGDRSSGGGYSGDRSGGGYGGDRS<br>GGGYGGDRGG<br>GYGGDRGGGYGGDRGGGYGGDRGGYGGDRGGGYGGDRG<br>GYGGDRGGYGG<br>DRGGYGGDRGGYGGDRSRGGYGGDRGGGSGYGGDRSGGY<br>GGDRSGGGYGG<br>DRGGGYGGDRGGYGGKMGGRNDYRNDQRNRPY<br>*(Red: TAF(1-208), black: TAF(209-592))* | |
| Tau | MAEPRQEFEVMEDHAGTYGLGDRKDQGGYTMHQDQEGDT<br>DAGLKESPLQT<br>PTEDGSEEPGSETSDAKSTPTAEDVTAPLVDEGAPGKQAAAQ<br>PHTEIPEGTTAE<br>EAGIGDTPSLEDEAAGHVTQARMVSKSKDGTGSDDKKAKG<br>ADGKTKIATPRGA<br>APPGQKGQANATRIPAKTPPAPKTPPSSGEPPKSGDRSGYSSP<br>GSPGTPGSRSR<br>TPSLPTPPTREPKKVAVVRTPPKSPSSAKSRLQTAPVPMPDLK<br>NVKSKIGSTENL<br>KHQPGGGKVQIINKKLDLSNVQSKCGSKDNIKHVPGGGSVQI<br>VYKPVDLSKVTS<br>KCGSLGNIHHKPGGGQVEVKSEKLDFKDRVQSKIGSLDNITH<br>VPGGGNKKIETH<br>KLTFRENAKAKTDHGAEIVYKSPVVSGDTSPRHLSNVSSTGSI<br>DMVDSPQLATLADEVSASLAKQGL<br>*(Red: Tau(1-150), black: Tau(151-441))* | Addgene Plasmid #16316 |
| Murine osteopontin (mOPN) | MRLAVICFCLFGIASSLPVKVTDSGSSEEKLYSLHPDPIATWL<br>VPDPSQKQNLLAP<br>QNAVSSEEKDDFKQETLPSNSNESHDHMDDDDDDDDDGD<br>HAESEDSVDSD<br>ESDESHHSDESDETVTASTQADTFTPIVPTVDVPNGRGDSLA<br>YGLRSKSRSFQV<br>SDEQYPDATDEDLTSHMKSGESKESLDVIPVAQLLSMPSDQD<br>NNGKGSHESSQ<br>LDEPSLETHRLEHSKESQESADQSDVIDSQASSKASLEHQSHK<br>FHSHKDKLVLDP<br>KCKEDDRYLKFRISHELESSSSEVN | Addgene Plasmid #106447 |
| DDX4 | MGDEDWEAEINPHMSSYVPIFEKDRYSGENGDNFNRTPASSS<br>EMDDGPSRRD<br>HFMKSGFASGRNFGNRDAGECNKRDNTSTMGGFGVGKSFG<br>NRGFSNSRFED | Addgene Plasmid #204408 |

| | | |
|---|---|---|
| | GDSSGFWRESSNDCEDNPTRNRGFSKRGGYRDGNNSEASGP YRRGGRGSFRG CRGGFGLGSPNNDLDPDECMQRTGGLFGSRRPVLSGTGNGD TSQSRSGSGS ERGGYKGLNEEVITGSGKNSWKSEAEGGES | |
| Alkaline phosphatase (ALP) | MPVLENRAAQGDITTPGGARRLTGDQTAALRDSLSDKPAKN IILLIGDGMGDS EITAARNYAEGAGGFFKGIDALPLTGQYTHYALNKKTGKPD YVTDSAASATAW STGVKTYNGALGVDIHEKDHPTILEMAKAAGLATGNVSTAE LQGATPAALVAH VTSRKCYGPSATSEKCPGNALEKGGKGSITEQLLNARADVTL GGGAKTFAETAT AGEWQGKTLREQAQTRGYQLVSDAASLNSVTEANQQKPLL GLFADGNMPVR WLGPKATYHGNIDKPAVTCTPNPQRNDSVPTLAQMTDKAIE LLSKNEKGFFLQ VEGASIDKQNHAANPCGQIGETVDLDEAVQRALEFAKKEGN TLVIVTADHAH ASQIVAPDTKAPGLTQALNTKDGAVMVMSYGNSEEDSQEHT GSQLRIAAYGP HAANVVGLTDQTDLFYTMKAALGLKVPSA | Addgene Plasmid #39265 |
| dCas9 | MDKKYSIGLAIGTNSVGWAVITDEYKVPSKKFKVLGNTDRH SIKKNLIGALLFDS GETAEATRLKRTARRRYTRRKNRICYLQEIFSNEMAKVDDSF FHRLEESFLVEED KKHERHPIFGNIVDEVAYHEKYPTIYHLRKKLVDSTDKADLR LIYLALAHMIKFRG HFLIEGDLNPDNSDVDKLFIQLVQTYNQLFEENPINASGVDA KAILSARLSKSRR LENLIAQLPGEKKNGLFGNLIALSLGLTPNFKSNFDLAEDAKL QLSKDTYDDDLD NLLAQIGDQYADLFLAAKNLSDAILLSDILRVNTEITKAPLSA SMIKRYDEHHQDL TLLKALVRQQLPEKYKEIFFDQSKNGYAGYIDGGASQEEFYK FIKPILEKMDGTE ELLVKLNREDLLRKQRTFDNGSIPHQIHLGELHAILRRQEDFY PFLKDNREKIEKIL TFRIPYYVGPLARGNSRFAWMTRKSEETITPWNFEEVVDKGA SAQSFIERMTN | NA |

| | | |
|---|---|---|
| | FDKNLPNEKVLPKHSLLYEYFTVYNELTKVKYVTEGMRKPA<br>FLSGEQKKAIVDL<br>LFKTNRKVTVKQLKEDYFKKIECFDSVEISGVEDRFNASLGT<br>YHDLLKIIKDKDFL<br>DNEENEDILEDIVLTLTLFEDREMIEERLKTYAHLFDDKVMK<br>QLKRRRYTGWGR<br>LSRKLINGIRDKQSGKTILDFLKSDGFANRNFMQLIHDDSLTF<br>KEDIQKAQVSG<br>QGDSLHEHIANLAGSPAIKKGILQTVKVVDELVKVMGRHKP<br>ENIVIEMARENQ<br>TTQKGQKNSRERMKRIEEGIKELGSQILKEHPVENTQLQNEK<br>LYLYYLQNGRD<br>MYVDQELDINRLSDYDVDAIVPQSFLKDDSIDNKVLTRSDKN<br>RGKSDNVPSEE<br>VVKKMKNYWRQLLNAKLITQRKFDNLTKAERGGLSELDKA<br>GFIKRQLVETRQI<br>TKHVAQILDSRMNTKYDENDKLIREVKVITLKSKLVSDFRKD<br>FQFYKVREINNY<br>HHAHDAYLNAVVGTALIKKYPKLESEFVYGDYKVYDVRKM<br>IAKSEQEIGKATA<br>KYFFYSNIMNFFKTEITLANGEIRKRPLIETNGETGEIVWDKG<br>RDFATVRKVLSM<br>PQVNIVKKTEVQTGGFSKESILPKRNSDKLIARKKDWDPKKY<br>GGFDSPTVAYSVL<br>VVAKVEKGKSKKLKSVKELLGITIMERSSFEKNPIDFLEAKG<br>YKEVKKDLIIKLPKYSL<br>FELENGRKRMLASAGELQKGNELALPSKYVNFLYLASHYEK<br>LKGSPEDNEQKQL<br>FVEQHKHYLDEIIEQISEFSKRVILADANLDKVLSAYNKHRD<br>KPIREQAENIIHLFT<br>LTNLGAPAAFKYFDTTIDRKRYTSTKEVLDATLIHQSITGLYE<br>TRIDLSQLGGD | |
| Carbonic anhydrase | MAHHWGYGKHNGPEHWHKDFPIAKGERQSPVDIDTHTAKY<br>DPSLKPLSVSYD<br>QATSLRILNNGHTFNVEFDDSQDKAVLKGGPLDGTYRLIQFH<br>FHWGSHDGQG<br>SEHTVDKKKYAAELHLVHWNTKYGDFGKAVQQPDGLAVL<br>GIFLKVGSANPGLQ<br>KVVDVLDSIKTKGKSADFTNFDPRGLLPESLDYWTYPGSLTT<br>PPLLECVTWIVLKE | NA |

| | | |
|---|---|---|
| | PISVSSEQVSKFRKLNFNGEGEPEEPMVDNWRPTQPLKNRQI KASFK | |
| YFP | MRKGEELFTGVVPILVELDGDVNGHKFSVSGEGEGDATYGK LTLKFICTTGKLPV PWPTLVTTFGYGLKCFARYPDHMKRHDFFKSAMPEGYVQE RTIFFKDDGNYKT RAEVKFEGDTLVNRIELKGIDFKEDGNILGHKLEYNYNSHNV YIMADKQKNGIKV NFKIRHNIEDGSVQLADHYQQNTPIGDGPVLLPDNHYLSYQS KLSKDPNEKRDH MVLLEFVTAAGITHGMDELYK | NA |
| sfGFP | MRKGEELFTGVVPILVELDGDVNGHKFSVRGEGEGDATNGK LTLKFICTTGKLP VPWPTLVTTLTYGVQCFARYPDHMKQHDFFKSAMPEGYVQ ERTISFKDDGTY KTRAEVKFEGDTLVNRIELKGIDFKEDGNILGHKLEYNFNSH NVYITADKQKNG IKANFKIRHNVEDGSVQLADHYQQNTPIGDGPVLLPDNHYLS TQSVLSKDPN EKRDHMVLLEFVTAAGITHGMDELYK | NA |
| CFP | MRKGEELFTGVVPILVELDGDVNGHKFSVSGEGEGDATYGK LTLKFICTTGKLP VPWPTLVTTLTWGVQCFSRYPDHMKRHDFFKSAMPEGYVQ ERTIFFKDDGNY KTRAEVKFEGDTLVNRIELKGIDFKEDGNILGHKLEYNYISH NVYITADKQKNGIK AHFKIRHNIEDGSVQLADHYQQNTPIGDGPVLLPDNHYLSTQ SKLSKDPNEKRD HMVLLEFVTAAGITHGMDELYK | NA |
| RFP (mCherry) | MVSKGEEDNMAIIKEFMRFKVHMEGSVNGHEFEIEGEGEGR PYEGTQTAKLK VTKGGPLPFAWDILSPQFMYGSKAYVKHPADIPDYLKLSFPE GFKWERVMNFE DGGVVTVTQDSSLQDGEFIYKVKLRGTNFPSDGPVMQKKTM GWEASSERMY PEDGALKGEIKQRLKLKDGGHYDAEVKTTYKAKKPVQLPG AYNVNIKLDITSHN EDYTIVEQYERAEGRHSTGGMDELYK | NA |
| $(PRM)_5$ | HMKGGSWGGSKKKKTAPTPPKRSGGSGGSGGSGGSKKKKT APTPPKRSGGS | Addgene Plasmid #175232 |

| | | |
|---|---|---|
| | GGSGGSGGSKKKKTAPTPPKRSGGSGGSGGSGGSKKKKTAP<br>TPPKRSGGSGG<br>SGGSGGSKKKKTAPTPPKRSGGSGS | |
| $(SUMO)_5$ | HMGGSWGGSMSEEKPKEGVKTENDHINLKVAGQDGSVVQF<br>KIKRHTPLSKL<br>MKAYSERQGLSMRQIRFRFDGQPINETDTPAQLEMEDEDTID<br>VFQQQTVVG<br>GSGGSGGSGGSMSEEKPKEGVKTENDHINLKVAGQDGSVVQ<br>FKIKRHTPLSK<br>LMKAYSERQGLSMRQIRFRFDGQPINETDTPAQLEMEDEDTI<br>DVFQQQTVV<br>GGSGGSGGSGGSMSEEKPKEGVKTENDHINLKVAGQDGSVV<br>QFKIKRHTPLS<br>KLMKAYSERQGLSMRQIRFRFDGQPINETDTPAQLEMEDEDT<br>IDVFQQQTVV<br>GGSGGSGGSGGSMSEEKPKEGVKTENDHINLKVAGQDGSVV<br>QFKIKRHTPLS<br>KLMKAYSERQGLSMRQIRFRFDGQPINETDTPAQLEMEDEDT<br>IDVFQQQTVV<br>GGSGGSGGSGGSMSEEKPKEGVKTENDHINLKVAGQDGSVV<br>QFKIKRHTPLS<br>KLMKAYSERQGLSMRQIRFRFDGQPINETDTPAQLEMEDEDT<br>IDVFQQQTVV<br>GGSGGS | Addgene Plasmid #126949 |
| $(SUMO)_{10}$ | GSGGSWGGSMSEEKPKEGVKTENDHINLKVAGQDGSVVQF<br>KIKRHTPLSKLM<br>KAYCERQGLSMRQIRFRFDGQPINETDTPAQLEMEDEDTIDV<br>FQQQTVVGGS<br>GGSGGSGGSMSEEKPKEGVKTENDHINLKVAGQDGSVVQFK<br>IKRHTPLSKLM<br>KAYCERQGLSMRQIRFRFDGQPINETDTPAQLEMEDEDTIDV<br>FQQQTVVGGS<br>GGSGGSGGSMSEEKPKEGVKTENDHINLKVAGQDGSVVQFK<br>IKRHTPLSKLM<br>KAYCERQGLSMRQIRFRFDGQPINETDTPAQLEMEDEDTIDV<br>FQQQTVVGGS<br>GGSGGSGGSMSEEKPKEGVKTENDHINLKVAGQDGSVVQFK<br>IKRHTPLSKLM<br>KAYCERQGLSMRQIRFRFDGQPINETDTPAQLEMEDEDTIDV<br>FQQQTVVGGS | Addgene Plasmid #126948 |

| | | |
|---|---|---|
| | GGSGGSGGSMSEEKPKEGVKTENDHINLKVAGQDGSVVQFK<br>IKRHTPLSKLM<br>KAYCERQGLSMRQIRFRFDGQPINETDTPAQLEMEDEDTIDV<br>FQQQTVVGGS<br>GGSGGSGGSMSEEKPKEGVKTENDHINLKVAGQDGSVVQFK<br>IKRHTPLSKLM<br>KAYCERQGLSMRQIRFRFDGQPINETDTPAQLEMEDEDTIDV<br>FQQQTVVGGS<br>GGSGGSGGSMSEEKPKEGVKTENDHINLKVAGQDGSVVQFK<br>IKRHTPLSKLM<br>KAYCERQGLSMRQIRFRFDGQPINETDTPAQLEMEDEDTIDV<br>FQQQTVVGGS<br>GGSGGSGGSMSEEKPKEGVKTENDHINLKVAGQDGSVVQFK<br>IKRHTPLSKLM<br>KAYCERQGLSMRQIRFRFDGQPINETDTPAQLEMEDEDTIDV<br>FQQQTVVGGS<br>GGSGGSGGSMSEEKPKEGVKTENDHINLKVAGQDGSVVQFK<br>IKRHTPLSKLM<br>KAYCERQGLSMRQIRFRFDGQPINETDTPAQLEMEDEDTIDV<br>FQQQTVVGGS<br>GGSGGSGGSMSEEKPKEGVKTENDHINLKVAGQDGSVVQFK<br>IKRHTPLSKLM<br>KAYCERQGLSMRQIRFRFDGQPINETDTPAQLEMEDEDTIDV<br>FQQQTVVGGS<br>CGGSLE | |
| $(SIM)_5$ | GSGGSWGGSKVDVIDLTIESSSDEEEDPPAKRGGSGGSGGSG<br>GSKVDVIDLTIE<br>SSSDEEEDPPAKRGGSGGSGGSGGSKVDVIDLTIESSSDEEED<br>PPAKRGGSGGS<br>GGSGGSKVDVIDLTIESSSDEEEDPPAKRGGSGGSGGSGGSK<br>VDVIDLTIESSSD<br>EEEDPPAKRGGSCGGSLE | Addgene Plasmid #126947 |
| $(SIM)_{10}$ | GSGGSWGGSKVDVIDLTIESSSDEEEDPPAKRGGSGGSGGSG<br>GSKVDVIDLTIE<br>SSSDEEEDPPAKRGGSGGSGGSGGSKVDVIDLTIESSSDEEED<br>PPAKRGGSGGS<br>GGSGGSKVDVIDLTIESSSDEEEDPPAKRGGSGGSGGSGGSK<br>VDVIDLTIESSSD<br>EEEDPPAKRGGSGGSGGSGGSKVDVIDLTIESSSDEEEDPPAK<br>RGGSGGSGGS | Addgene Plasmid #126946 |

| | | |
|---|---|---|
| | GGSKVDVIDLTIESSSDEEEDPPAKRGGSGGSGGSGGSKVDVI DLTIESSSDEEE DPPAKRGGSGGSGGSGGSKVDVIDLTIESSSDEEEDPPAKRG GSGGSGGSGGS KVDVIDLTIESSSDEEEDPPAKRGGSCGGSLE | |
| $(SH3)_3$ | MDLNMPAYVKFNYMAEREDELSLIKGTKVIVMEKSSDGWW RGSYNGQVGW FPSNYVTEEGDSPLASGAGGSEGGGSEGGTSGATDLNMPAY VKFNYMAERED ELSLIKGTKVIVMEKSSDGWWRGSYNGQVGWFPSNYVTEEG DSPLASGAGGS EGGGSEGGTSGATDLNMPAYVKFNYMAEREDELSLIKGTKV IVMEKSSDGWW RGSYNGQVGWFPSNYVTEEGDSPLGGGS | Addgene Plasmid #175233 |
| ELP(S96) | MGVPGSGVPGSGVPGSGVPGSGVPGSGVPGSGVPGSGVPGS GVPGSGVPGS GVPGSGVPGSGVPGSGVPGSGVPGSGVPGSGVPGSGVPGSG VPGSGVPGSGV PGSGVPGSGVPGSGVPGSGVPGSGVPGSGVPGSGVPGSGVPG SGVPGSGVPG SGVPGSGVPGSGVPGSGVPGSGVPGSGVPGSGVPGSGVPGSG VPGSGVPGSG VPGSGVPGSGVPGSGVPGSGVPGSGVPGSGVPGSGVPGSGVP GSGVPGSGVP GSGVPGSGVPGSGVPGSGVPGSGVPGSGVPGSGVPGSGVPGS GVPGSGVPGS GVPGSGVPGSGVPGSGVPGSGVPGSGVPGSGVPGSGVPGSG VPGSGVPGSGV PGSGVPGSGVPGSGVPGSGVPGSGVPGSGVPGSGVPGSGVPG SGVPGSGVPG SGVPGSGVPGSGVPGSGVPGSGVPGSGVPGSGVPGSGVPGSG VPGSGVPGSG VPGSGVPGSGVPGSGY | Addgene Plasmid # 68393 |
| ELP(S48I 48) | MGVPGSGVPGSGVPGSGVPGSGVPGSGVPGSGVPGSGVPGS GVPGSGVPGS GVPGSGVPGSGVPGSGVPGSGVPGSGVPGSGVPGSGVPGSG VPGSGVPGSGV PGSGVPGSGVPGSGVPGSGVPGSGVPGSGVPGSGVPGSGVPG SGVPGSGVPG SGVPGSGVPGSGVPGSGVPGSGVPGSGVPGSGVPGSGVPGSG VPGSGVPGS | Addgene Plasmid # 68394 |

| | | |
|---|---|---|
| | GVPGSGVPGSGVPGSGVPGSGVPGSGVPGSGVPGSGVPGIGV PGIGVPGIGV PGIGVPGIGVPGIGVPGIGVPGIGVPGIGVPGIGVPGIGVPGIGV PGIGVPGIGV PGIGVPGIGVPGIGVPGIGVPGIGVPGIGVPGIGVPGIGVPGIGV PGIGVPGIGV PGIGVPGIGVPGIGVPGIGVPGIGVPGIGVPGIGVPGIGVPGIGV PGIGVPGIGV PGIGVPGIGVPGIGVPGIGVPGIGVPGIGVPGIGVPGIGVPGIGV PGIGVPGIGV PGIGY | |
| α-Synuclein | MDVFMKGLSKAKEGVVAAAEKTKQGVAEAAGKTKEGVLY VGSKTKEGVVHG VTTVAEKTKEQVTNVGGAVVTGVTAVAQKTVEGAGNIAAA TGFVKKDQMGK GEEGYPQEGILEDMPVDPGSEAYEMPSEEGYQDYEPEA | Addgene Plasmid # 166671 |
| hnRNAP A1LCD | MASASSSQRGRSGSGNFGGGRGGGFGGNDNFGRGGNFSGR GGFGGSRGGGG YGGSGDGYNGFGNDGSNFGGGGSYNDFGNYNNQSSNFGPM KGGNFGGRSSG PYGGGGQYFAKPRNQGGYGGSSSSSSYGSGRRF | Addgene Plasmid #204405 |
| RGG-RGG | MESNQSNNGGSGNAALNRGGRYVPPHLRGGDGGAAAAASA GGDDRRGGAG GGGYRRGGGNSGGGGGGGYDRGYNDNRDDRDNRGGSGGY GRDRNYEDRGY NGGGGGGGNRGYNNNRGGGGGGYNRQDRGDGGSSNFSRG GYNNRDEGSD NRGSGRSYNNDRRDNGGDGEFGKLMESNQSNNGGSGNAAL NRGGRYVPPHL RGGDGGAAAAASAGGDDRRGGAGGGGYRRGGGNSGGGGG GGYDRGYND NRDDRDNRGGSGGYGRDRNYEDRGYNGGGGGGGNRGYNN NRGGGGGGY NRQDRGDGGSSNFSRGGYNNRDEGSDNRGSGRSYNNDRRD NGGDG | Addgene Plasmid # 124941 |
| RGG | MESNQSNNGGSGNAALNRGGRYVPPHLRGGDGGAAAAASA GGDDRRGGA GGGGYRRGGGNSGGGGGGGYDRGYNDNRDDRDNRGGSGG YGRDRNYEDR GYNGGGGGGGNRGYNNNRGGGGGGYNRQDRGDGGSSNFS RGGYNNRDE | |

| | | |
|---|---|---|
| | GSDNRGSGRSYNNDRRDNGGDG | |
| FUSLC | MASNDYTQQATQSYGAYPTQPGQGYSQQSSQPYGQQSYSG YSQSTDTSGYG QSSYSSYGQSQNTGYGTQSTPQGYGSTGGYGSSQSSQSSYGQ QSSYPGYGQQ PAPSSTSGSYGSSSQSSSYGQPQSGSYSQQPSYGGQQQSYGQ QQSYNPPQGYG QQNQYNS | Addgene Plasmid # 127192) |
| $(XGZG)_2$-X | 1. FGFGFGFGFGFGFGFGF<br>2. IGIGIGIGIGIGIGIGI<br>3. QGQGQGQGQGQGQGQGQ<br>4. RGRGRGRGRGRGRGRGR<br>5. SGSGSGSGSGSGSGSGS<br>6. WGWGWGWGWGWGWGWGW<br>7. YGYGYGYGYGYGYGYGY<br>8. DGDGDGDGDGDGDGDGD<br>9. RGDGRGDGRGDGRGDGR | Extension primers of PCR |

Note: NA means the protein genes were already provided in lab. And all protein genes were confirmed via sequencing.

**Table S5. Plasmids used in this work**

| Plasmids | Description | Antibiotic resistance | Source |
|---|---|---|---|
| pBbB8k-csg-nb112 | Arabinose-inducible csgBACEFG operon for curli fiber synthesis in which curli subunit CsgA has been fused to a nanobody targeting the spike protein of SARS-CoV-2 via a flexible linker. | Kanamycin | Ref.[2] |
| pBbB8k-ΔBAC_CsgEFG | Traceless deletion of CsgB, CsgA and CsgC. | Kanamycin | This work |
| pBbB8k-ΔBAC_CsgF-fusion protein tag | Traceless deletion of CsgB, CsgA and CsgC.<br>CsgF fused with additional protein tag (including His tag, FLAG tag and Spy tag) at the C-terminal. | Kanamycin | This work |
| pBbB8k-ΔBAC_CsgF-fusion protein sequences containing His tag | Traceless deletion of CsgB, CsgA and CsgC.<br>CsgF fused with additional protein segment or sequences (including p53(1-393), p53(1-93), TAF(1-592), TAF(1-234), Tau(1-441), Tau(1-150), Murine osteopontin (mOPN), DDX4, dCas9, ALP, Carbonic anhydrase, YFP, sfGFP, CFP, $(PRM)_5$, $(SUMO)_5$, $(SUMO)_{10}$, CsgA-FUSLC, $(SIM)_5$, $(SIM)_{10}$, $(SH3)_3$, ELP(S96), ELP(S48I48), α-Synuclein, hnRNAPA1LCD, RGG-RGG, RGG, FUSLC) containing His tag, as shown in Table S4. | Kanamycin | This work |
| pBbB8k-ΔBAC_CsgF-FUSLC-FLAG tag | Traceless deletion of CsgB, CsgA and CsgC.<br>CsgF fused with FUSLC-His tag at the C-terminal. | Kanamycin | This work |
| pBbB8k-ΔBAC_CsgF-FUSLC-FLAG tag_CsgF-RGG-His tag | Traceless deletion of CsgB, CsgA and CsgC.<br>CsgF fused with FUSLC-His FLAG tag at the C-terminal, and additional involvement of CsgF-RGG-His tag. | Kanamycin | This work |
| pBbB8k-ΔBAC_CsgF- | Traceless deletion of CsgB, CsgA and CsgC. | Kanamycin | This work |

| | | | |
|---|---|---|---|
| (XGZG)$_2$-X-His tag | CsgF fused with (XGZG)$_2$-X-His tag at the C-terminal, in which X, Z could represent natural amino acids like Q, S, Y, W, F, I, D, R. | | |
| pBbB8k-ΔBA-CsgF-His tag | Traceless deletion of CsgB, CsgA. CsgF fused with His tag. | Kanamycin | This work |
| pBbB8k-ΔBAG_CsgF-His tag | Traceless deletion of CsgB, CsgA, CsgG.<br>CsgF fused with His tag. | Kanamycin | This work |
| pBbB8k-ΔBACG _CsgF-His tag | Traceless deletion of CsgB, CsgA, CsgC and CsgG. CsgF fused with His tag. | Kanamycin | This work |
| pBbB8k-ΔBAC_CFP_CsgF | Traceless deletion of CsgB, CsgA, and CsgC. Insertion of intracellular expression of CFP. | Kanamycin | This work |
| pBbB8k-ΔBAC_RFP_CsgF | Traceless deletion of CsgB, CsgA, and CsgC. Insertion of intracellular expression of RFP. | Kanamycin | This work |
| pBbB8k-ΔBAC_CFP_CsgF-FUSLC | Traceless deletion of CsgB, CsgA, and CsgC. CsgF fused with FUSLC. Insertion of intracellular expression of CFP. | Kanamycin | This work |
| pBbB8k-ΔBAC_RFP_CsgF-FUSLC | Traceless deletion of CsgB, CsgA, and CsgC. CsgF fused with FUSLC. Insertion of intracellular expression of RFP. | Kanamycin | This work |
| pBbB8k-ΔBAC_YFP_CsgF-FUSLC | Traceless deletion of CsgB, CsgA, and CsgC. CsgF fused with FUSLC. Insertion of intracellular expression of YFP. | Kanamycin | This work |
| pBbB8k-ΔBAC_CFP_CsgF-RGG | Traceless deletion of CsgB, CsgA, and CsgC. CsgF fused with RGG. Insertion of intracellular expression of CFP. | Kanamycin | This work |
| pBbB8k-ΔBAC_RFP_CsgF-RGG | Traceless deletion of CsgB, CsgA, and CsgC. CsgF fused with RGG. Insertion of intracellular expression of RFP. | Kanamycin | This work |
| pBbB8k-ΔBAC_YFP_CsgF-FUSLC_CsgF-RGG | Traceless deletion of CsgB, CsgA, and CsgC. CsgF fused with FUSLC and additional insertion of CsgF fused with RGG. Insert of intracellular expression of YFP. | Kanamycin | This work |

| | | | |
|---|---|---|---|
| pBbB8k-ΔBAC_CFP_CsgF-ELP(S96) | Traceless deletion of CsgB, CsgA, and CsgC. CsgF fused with ELP(S96). Insertion of intracellular expression of CFP. | Kanamycin | This work |
| pBbB8k-ΔBAC_CFP_CsgF-ELP(S48I48) | Traceless deletion of CsgB, CsgA, and CsgC. CsgF fused with ELP(S96). Insertion of intracellular expression of CFP. | Kanamycin | This work |
| pBbB8k-ΔBAC_Sec-N22-FUSLC_CsgF-FUSLC | Traceless deletion of CsgB, CsgA, and CsgC. CsgF fused with FUSLC. Additional insertion of FUSLC with secreted tag. | Kanamycin | This work |
| pBbB8k-ΔBAC_Sec-N22-$(SUMO)_5$ | Traceless deletion of CsgB, CsgA, and CsgC. Insertion of $(SUMO)_5$ with secreted tag. | Kanamycin | This work |
| pBbB8k-ΔBAC_Sec-N22-RGG_CsgF-RGG | Traceless deletion of CsgB, CsgA, and CsgC. CsgF fused with RGG. Additional insertion of RGG with secreted tag. | Kanamycin | This work |
| pBbB8k-ΔB_CsgA-FUSLC_CsgF-FUSLC | Traceless deletion of CsgB. CsgF fused with FUSLC. Additional fusion of FUSLC with CsgA at the C-terminal. | Kanamycin | This work |
| pBbB8k-ΔB_CsgA-RGG_CsgF-RGG | Traceless deletion of CsgB. CsgF fused with RGG. Additional fusion of RGG with CsgA at the C-terminal. | Kanamycin | This work |
| pBbB8k-ΔB_CsgA-RGG_CsgF-RGG | Traceless deletion of CsgB. CsgF fused with RGG. Additional fusion of RGG with CsgA at the C-terminal. | Kanamycin | This work |
| pET21d-CsgF-His tag | Expression of CsgF-His tag and purification. | Ampicillin | This work |

**Table S6. Strains used in this work**

| Strains | Description | Source |
|---|---|---|
| PQN4 | An *E. coli* cell strain derived from LSR10 (MC4100, ΔcsgA, λ(DE3), CamR) with deletion of curli operon (ΔcsgBACEFG).<br>In this work, all PQN4 strains transformed with pBbB8k backbone-based plasmids, as shown Table S5. | Ref.[4] |
| BL21(DE3) | A commercial *E. coli* strain for protein expression. | New England Biolabs |
